# Alternative splicing links sex to neutrophil inflammatory vulnerability

**DOI:** 10.64898/2026.11.24.769038

**Authors:** Mayumi Mori, Chiaki Abe, Satoko Sakurai, Taro Tsujimura, Shohei Murakami, Hozumi Motohashi, Takuya Yamamoto, Yo-ichi Nabeshima

**Author notes:** Correspondence to: Mayumi Mori, Department of Aging Science and Medicine, Graduate School of Medicine, Kyoto University, Shogoin Kawahara-cho 53, Sakyo-ku, 606-8607 Kyoto, Japan.

## Abstract

Mammalian females generally live longer than males, yet the immunological drivers governing sex-biased inflammatory mortality remain unclear. Neutrophils, among conserved immune cells across species, exhibit marked sex differences in inflammatory and age-related diseases. We identified conserved male-biased CD11b expression in resting neutrophils of wild-type mice, *Rag2* KO mice, and in human cohorts. Male neutrophils selectively displayed CD11b-dependent transendothelial migration. During tissue inflammation in our mouse model deficient in *α-Klotho* (*αKl*) combined with *Rag2* KO lymphocyte depletion (dKO), males showed higher mortality and more severe pulmonary pathology correlated with increased infiltrating neutrophils in the lung. The neutrophil accumulation directly accounted for heightened IL-1β expression in males despite equivalent per-cell cytokine expression across sexes. Transcriptomic analysis of neutrophils revealed profound sex-biased alternative splicing enriched in Toll-like receptor (TLR) cascades, alongside hormone-responsive regulation of integrin β2 (*Itgb2*) isoform usage, implicating alternative splicing in neutrophils represents male vulnerability to tissue inflammation.

**In Brief:** Mori et al. reveal that male susceptibility to inflammatory tissue damage is pre-programmed in resting neutrophils through sex-biased alternative RNA splicing of Toll-like receptor pathways-related genes, including *Itgb2*. The essential integrin partner for transendothelial migration, CD11b, is basally elevated in male neutrophils, which accounts for increased pulmonary neutrophil infiltration and neutrophil-derived IL-1β accumulation, dictating male-biased emphysema and mortality.

**Highlights:**

- Male neutrophils without priming possess an elevated surface CD11b expression in both mice and human
- Transendothelial migration of male but not female neutrophils is CD11b-dependent
- Emphysemic pulmonary pathology and mortality together with lung neutrophil infiltration are enhanced in male mice
- Sex-biased alternative splicing preconfigures Toll-like receptor cascades including Itgb2 isoforms

## Introduction

Across multiple species, biological sex is a fundamental determinant of immunity, disease susceptibility, and lifespan. Mammalian females generally exhibit stronger immune responses and lower mortality rates from many infectious and inflammatory conditions, whereas males often show greater susceptibility to tissue damage and inflammatory pathology.^1−3^ While adaptive immune responses contribute significantly to sex-biased autoimmunity and pathogen clearance, growing evidence indicates that innate immune cell-intrinsic dimorphisms represent a fundamental yet understudied determinant of differential inflammatory injury and tissue deterioration.^3,4^

Neutrophils, the primary cellular responders of the innate immune system, undergo rapid bone marrow mobilization, intravascular adhesion, and transendothelial migration into peripheral tissues upon inflammatory challenges.^5,6^ Recent studies have highlighted neutrophils as active regulators of sterile inflammation, tissue remodeling, and aging-associated pathology, rather than merely terminal antimicrobial effector cells.^7−10^ However, sexual dimorphism in neutrophils regarding adhesion mechanics and transcriptional or post-transcriptional molecular processing associated with systemic disease susceptibility has been poorly investigated.

Neutrophil trafficking is critically controlled by β2 integrins, particularly the Mac-1 complex, which is composed of CD11b (*Itgam*) and CD18 (*Itgb2*).^11,12^ Beyond its established role in firm adhesion to activated endothelium, CD11b regulates neutrophil recruitment, transendothelial migration, tissue retention, and inflammatory signaling.^13^ Elevated surface exposure of CD11b is widely used as a marker of neutrophil activation and mobilization.^14,15^ In the lung, the pulmonary capillary network serves as a major physiological reservoir for marginated neutrophils and is highly dependent on integrin-mediated adhesive interactions.^16^ Neutrophil recruitment and persistence within the inflamed lung can amplify local cytokine production, particularly IL-1β-dependent inflammatory circuits,^17^ thereby contributing to chronic tissue damage.^18,19^ Despite its central role in neutrophil biology, how integrin adhesion exhibits physiological sexual dimorphism and contributes to sex-dependent inflammatory susceptibility remains largely unknown.

Emerging transcriptomic studies suggest that alternative splicing represents a major source of functional diversity within terminally differentiated immune populations.^20−22^ Accumulating evidence suggests that alternative isoforms can modulate receptor signaling, adhesion, and inflammatory responsiveness in lymphocytes.^23,24^ However, although a recent multiomics study suggested that sex-dependent molecular programs exist in neutrophils,^25^ sex-dependent splicing programs associated with neutrophil phenotypes have not been systematically explored.

During investigations of inflammatory pathology in *α-Klotho*-deficient (*αKl* KO) mice, we identified an unexpected sex difference in neutrophil phenotype. *αKl* KO mice represent a classic, widely recognized progeria model characterized by accelerated aging, severe tissue pathology, and shortened lifespan.^26,27^ The aging-like pathology such as pulmonary emphysema, ectopic tissue calcification, and osteoporosis in *αKl* KO mice is associated with dysregulated mineral metabolism via profound hypervitaminosis D.^28^ Among the organs affected by *αKl* deficiency, the lung is particularly susceptible to structural degeneration, exhibiting emphysematous remodeling, alveolar destruction, and pathological mineral deposition.^29^ In our newly generated lymphocyte-deficient double-knockout model by crossing *αKl* κΟ mice with *Rag2* KO animals (dKO), aimed to dissecting the relative contributions of adaptive versus innate immune mechanisms to this pulmonary phenotype, male-biased mortality was expanded regardless of depleted adaptive lymphocytes and irrespective of circulating vitamin D. Additionally, we observed a robust male-biased increase in neutrophil surface CD11b expression, which was more broadly detected in wild-type mice and human datasets.

Here we hypothesized that sexual dimorphism in neutrophils may contribute to male-biased inflammatory susceptibility and explored the molecular mechanisms underlying these differences via global alternative mRNA splicing (AS) analysis across public and our original datasets in addition to conventional gene expression analysis.

## Results

### Lymphocyte deficiency expands male-biased mortality and pulmonary pathology

Although systemic inflammatory susceptibility has been reported in *α-Klotho* (*αKl*)-deficient mice,^30^ organ-or tissue-specific pathology such as pulmonary emphysema has been poorly addressed immunologically. To investigate whether the adaptive immune system modulates the pathological phenotypes driven by *αKl* deficiency, we crossed *Rag2* KO mice with *αKl* KO mice to generate lymphocyte-deficient double-knockout (dKO) animals. Reassessment of our previous data^31^ by Kaplan‒Meier survival analysis revealed that male *αKl* KO mice exhibited a modest trend toward male mortality (**Figure 1A**). Notably, this sex-dimorphic survival pattern was clearly observed in dKO mice by both log-rank and Gehan‒Wilcoxon tests.

**Figure 1.**
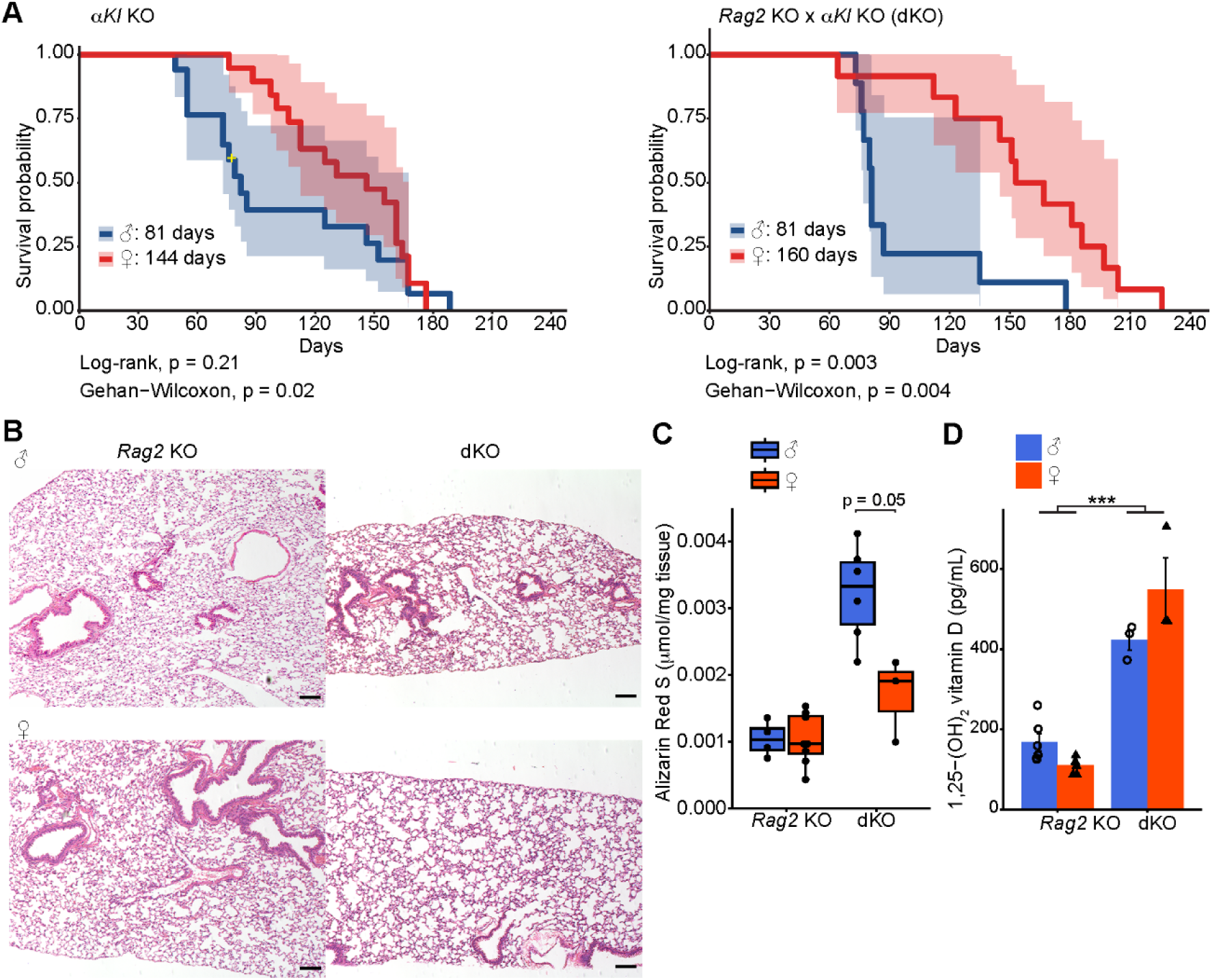
Lymphocyte deficiency expands male-biased mortality and pulmonary calcification in *αKl*-deficient mice, independently of systemic vitamin D levels. **(A)** Kaplan–Meier survival curves of male (blue) and female (red) *αKl* KO (left, *n* = 17 male, 19 female) and *Rag2* KO x *αKl* KO (dKO) mice (right, *n* = 9 male, 12 female). Shaded bands represent 95% confidence intervals. Insets display median survival in days. Log-rank (χ^2^ = 1.6 for *αKl* KO, 9 for dKO) and Gehan–Wilcoxon tests (χ^2^ = 5.5 for *αKl* KO, 8.2 for dKO) are shown below. **(B)** Representative hematoxylin and eosin (H&E) stained lung sections from male (top row) and female (bottom row) *Rag2* KO and dKO mice. Scale bars = 100 μm. **(C)** Quantification of pulmonary calcification measured by Alizarin Red S assay (μmol/mg tissue) in male (blue) and female (red) *Rag2* KO and dKO mice (*n* = 3–8 per group). Box plots display the median, interquartile range, and individual biological replicates. Statistical evaluation by two-way ANOVA followed by post-hoc pairwise comparisons with Benjamin-Hochberg adjustment. **(D)** Serum levels of 1,25-(OH)_2_ vitamin D (pg/mL) measured in male (blue) and female (red) *Rag2* KO and dKO mice (*n* = 3–7 per group). Data are presented as mean ± SEM with individual data points. Statistical significance was determined by two-way ANOVA (***: *p* < 0.001 for dKO vs. *Rag2* KO main effect; no significant sex difference within dKO).

Consistent with the known phenotypes of *αKl* KO mice,^27,29^ histological examination of lung sections revealed extensive pulmonary emphysema in dKO mice (**Figure 1B**). Likewise, compared to *Rag2* KO controls, dKO mice presented growth retardation accompanied by body weight loss (**Figure S1A**). Quantitative assessment by the Alizarin Red S assay on pulmonary homogenates confirmed a marked increase in pulmonary calcium deposition in male dKO mice and, to a lesser extent, in female dKO mice (**Figure 1C**). The severity of lung lesions paralleled the survival data, supporting the association between enhanced pulmonary pathology and reduced lifespan.

Given that hypervitaminosis D is a primary driver of systemic tissue calcification in *αKl*-deficient models,^28^ we assessed circulating concentrations of 1,25-(OH)₂ vitamin D. Serum vitamin D levels were significantly increased in dKO mice compared with *Rag2* KO controls, consistent with the known metabolic consequences of *αKl* deficiency (**Figure 1D**). However, no significant differences were detected between male and female dKO mice despite the marked sex differences in survival and pulmonary calcification. Moreover, systemic phosphorus/calcium homeostasis appears to be comparable between sexes (**Figures S1B and S1C**). These findings suggest that the exacerbated mortality and lung calcification observed in male dKO mice occur independently of vitamin D-mediated mineral metabolism.

Collectively, these findings indicate that in the absence of T and B lymphocytes, innate immune cells may exacerbate the male-biased pulmonary lesions associated with *αKl* deficiency, without affecting systemic vitamin D levels.

### Male-biased elevation of neutrophil CD11b expression is a conserved trait across mouse genotypes and human cohorts

To determine whether sexual dimorphism of innate immune cells reflects a general biological property rather than a disease-specific consequence of *αKl* deficiency, we first examined myeloid development in the bone marrow of WT, *αKl* KO, *Rag2* KO, and dKO mice. Flow cytometric analysis revealed no consistent sex-dependent differences in the frequencies of monocyte or neutrophil progenitors across genotypes (**Figures 2A, S2A, and S2B**). Furthermore, while mature CD101^+^ neutrophils were reduced in both *αKl* KO and dKO bone marrows, there were no sex-specific shifts (**Figure 2A**). Similarly, with physiological aging in both WT and *Rag2* KO mice, mature neutrophils were reduced, while neutrophil progenitors were sustained (**Figure 2B**). This suggests that the decrease in bone marrow neutrophils in *αKl* KO and dKO mice is associated with their progeric features, independent of sex-biased pulmonary phenotypes.

**Figure 2.**
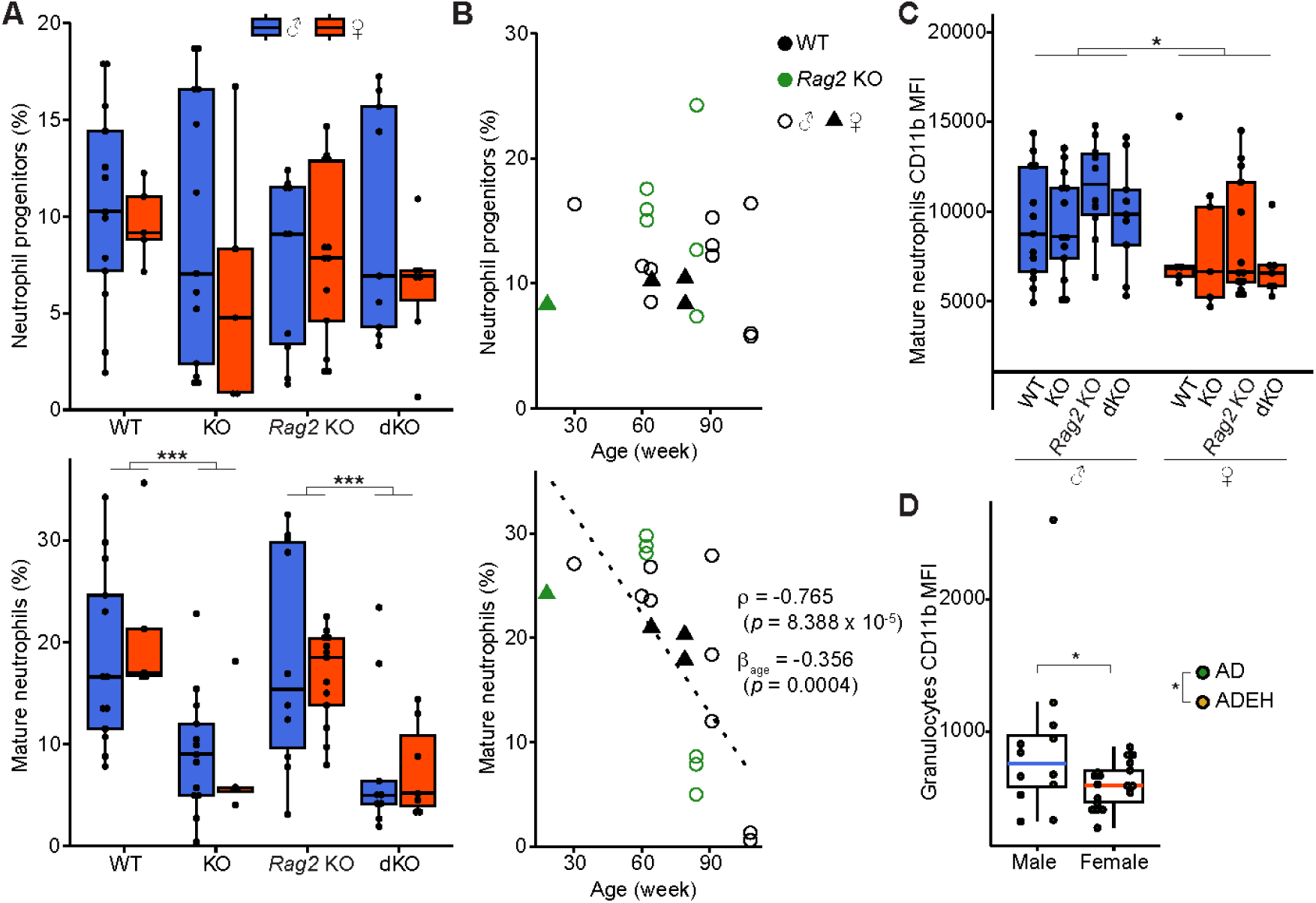
Male-biased elevation of neutrophil CD11b expression is a conserved trait across mouse genotypes and human cohorts. **(A)** Flow cytometric quantification of neutrophil progenitor frequencies (top) and CD101^+^ mature neutrophil frequencies (bottom) in the bone marrow of male (blue) and female (red) WT, *αKl* KO, *Rag2* KO, and dKO mice (*n* = 5–13 per group). Box plots display the median, interquartile range, and individual biological replicates. **(B)** Age-dependent frequency dynamics of neutrophil progenitors (top) and mature neutrophils (bottom) plotted against age (weeks) in male (open circles) and female (filled triangles) WT (black) and *Rag2* KO (green) mice aged > 12 weeks. Statistical evaluation demonstrated an age-dependent decline in mature neutrophils, as assessed by Spearman’s rank correlation (ρ = -0.765, *p* = 8,388 × 10^-5^) and linear regression (β_age_ = -0.356, *p* = 0.0004). **(C)** Quantification of surface CD11b mean fluorescence intensity (MFI) on bone marrow neutrophils from male and female mice across WT, *αKl* KO, *Rag2* KO, and dKO genotypes (*n* = 5–13 per group). Statistical evaluation by two-way ANOVA demonstrated a significant main effect of sex on CD11b MFI, regardless of genotype (*: *p* < 0.05). **(D)** Surface CD11b MFI on circulating granulocytes isolated from male (blue) and female (red) patient cohorts with atopic dermatitis (AD, green circles) or AD with eczema herpeticum (ADEH, yellow circles). Statistical evaluation by two-way ANOVA followed by post-hoc pairwise comparisons with Benjamin-Hochberg adjustment (*: *p* < 0.05).

We then evaluated the expression of integrin CD11b, a marker associated with neutrophil activation and functional responsiveness. In contrast to the minimal sex effects observed on neutrophil frequencies, surface CD11b expression on bone marrow

neutrophils was consistently elevated in males relative to females (**Figure 2C**). Notably, the male-biased increase in CD11b expression was observed in WT, *αKl* KO, *Rag2* KO, and dKO mice, indicating that heightened CD11b expression represents a conserved feature of male neutrophils rather than a secondary consequence of lymphocyte deficiency or pathology associated with *α-Klotho* loss.

To test whether this sex disparity translates to human biology, we analyzed surface CD11b mean fluorescence intensity (MFI) in circulating granulocytes using data from human patient cohorts with atopic dermatitis (AD) and AD complicated by eczema herpeticum (ADEH). Similar to the findings in mice, male subjects exhibited higher CD11b expression on granulocytes but not on non-granulocytes than female subjects across disease categories (**Figures 2D and S2C**). Taken together, these data demonstrate that elevated neutrophil CD11b expression represents a conserved male sex-dependent phenotype across mammalian genetic backgrounds. In contrast, neutrophil developmental frequencies showed minimal sex-specific variation, implicating qualitative differences in the neutrophil state, rather than quantitative differences in neutrophil production, as a potential contributor to the heightened male susceptibility observed in *αKl*-deficient mice.

### Transendothelial migration and pulmonary infiltration of neutrophils are associated with male-biased CD11b expression

We next assessed whether CD11b mediates neutrophil extravasation in a sex-dependent manner by evaluating the functional necessity of CD11b using an *in vitro* endothelial transmigration assay.^32,33^ Primary bone marrow neutrophils isolated from male and female WT mice were treated with either an anti-CD11b neutralizing antibody (αCD11b) or an isotype control antibody (Ctrl) and subjected to chemoattractant-driven transmigration across a bEnd.3 endothelial monolayer (**Figure 3A**). CD11b blockade reduced the transmigration of male WT neutrophils relative to that of control-treated cells in two out of three experiments. In contrast, female neutrophil migration was unaffected by CD11b blockade in all three experiments, indicating negligible sensitivity to the inhibition of integrin-mediated cell-to-cell contact (**Figure 3A**). These results demonstrate that male neutrophils rely more heavily on CD11b-dependent mechanisms for transendothelial migration.

**Figure 3.**
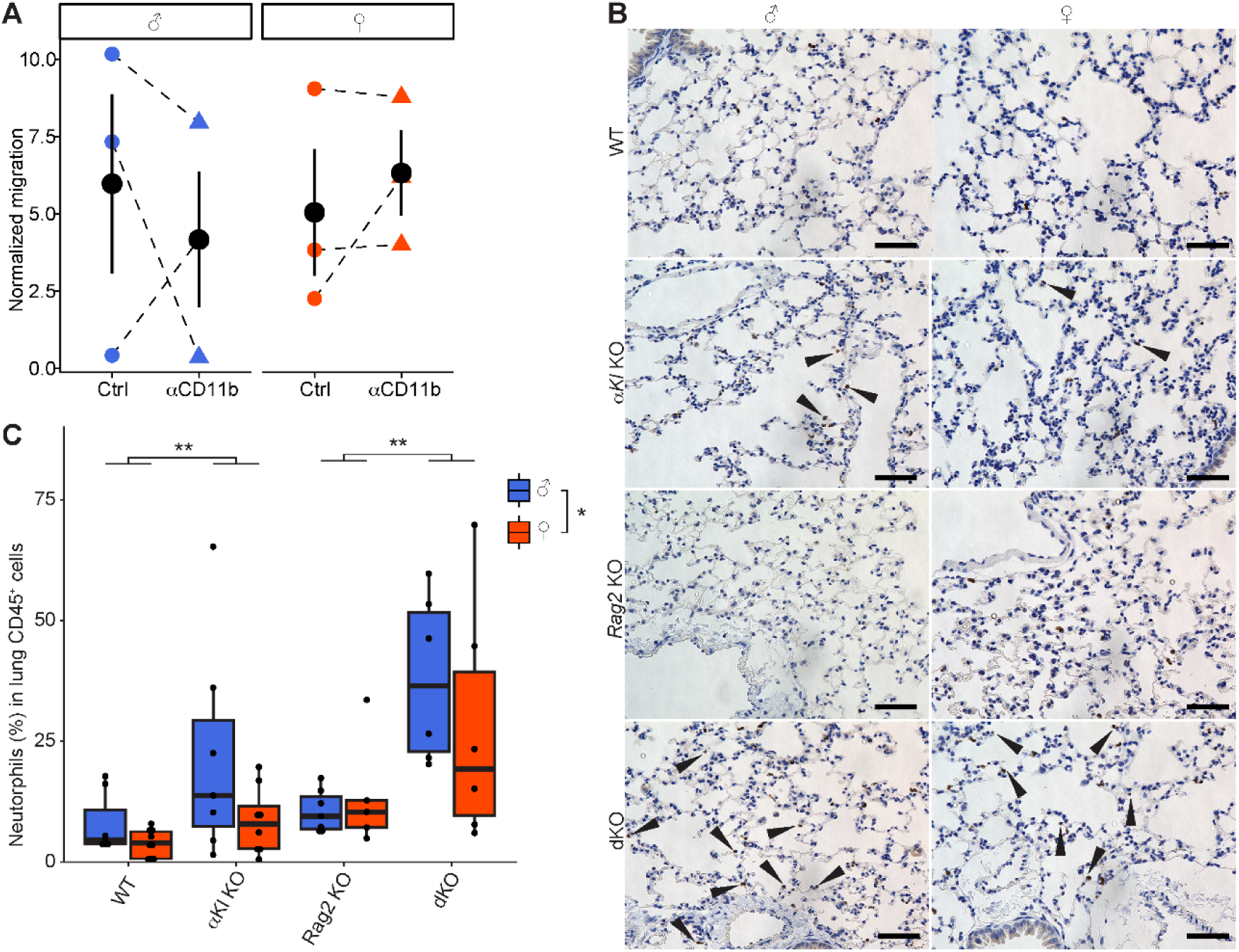
CD11b-dependent male neutrophil transmigration *in vitro* and increased pulmonary infiltration *in vivo*. **(A)** *In vitro* transmigration assay of bone marrow neutrophils isolated from wild-type (WT) male (blue, left) and female (red, right) mice, treated with control isotype IgG (Ctrl) or anti-CD11b neutralizing antibody (αCD11b). The number of Ly6G^+^ cells in the bottom wells migrated toward CXCL2 through the bEnd.3 layers on the upper wells was normalized to gravity-dependent backgrounds without chemoattractant. The dashed lines connect individual paired experimental replicates; black dots and error bars represent mean ± SEM (*n* = 3 per sex, both biologically and technically independent). **(B)** Representative immunohistochemical (IHC) detection of Ly6G^+^ neutrophils in the lung parenchyma of male (left column) and female (right column) WT, *αKl* KO, *Rag2* KO, and dKO mice. Black arrowheads indicate infiltrated Ly6G^+^ neutrophils. Scale bars = 100 μm. **(C)** Flow cytometric quantification of neutrophils (Ly6G^+^CD11b^+^CD11c^-^) expressed as a percentage of total lung CD45^+^ leukocytes in male (blue) and female (red) mice across WT, *αKl* KO, *Rag2* KO, and dKO genotypes (*n* = 5–9 per group). Box plots display the median, interquartile range, and individual biological replicates. Statistical evaluation by two-way ANOVA followed by post-hoc pairwise comparisons with Benjamin-Hochberg adjustment (**: *p* < 0.01 for genotype main effect; *: *p* < 0.05 for sex difference).

We further investigated whether these sex-specific differences in neutrophil migration translate to pulmonary neutrophil infiltration *in vivo*. Immunohistochemical staining for Ly6G revealed sparse neutrophil presence in WT or *Rag2* KO lungs of both sexes, whereas markedly increased numbers of Ly6G^+^ neutrophils were observed in *αKl* KO and dKO lungs. (**Figure 3B**). Neutrophil infiltration, identified as the extravascular deposition of Ly6G^+^ cells, was notably more pronounced in male mutant mice than in their female counterparts, with such accumulation remarkably detected in male dKO lungs (**Figure 3B**).

To precisely quantify the recruited population, we performed flow cytometry of lung CD45⁺ leukocytes to determine the proportion of Ly6G⁺CD11b⁺ neutrophils. Consistent with the histological findings, *αKl* KO and dKO strains exhibited elevated neutrophil proportions relative to WT and *Rag2* KO controls, respectively (**Figure 3C**). Furthermore, a significant sex-dependent difference was detected, with the most pronounced accumulation detected in the lungs of male dKO mice (**Figure 3C**).

Together, these findings indicate that CD11b drives sex-dimorphic neutrophil transmigration, mediating increased pulmonary neutrophilic inflammation in males. Analysis of other lung leukocyte populations, including eosinophils, macrophages, and lymphocytes, revealed no significant differences between *αKl*-deficient and control mice (**Figures S3A and S3B**). Similarly, leukocyte profiling in the kidneys demonstrated no notable alterations across genotypes, indicating that the neutrophilic inflammation observed was specific to the lung tissue (**Figures S3C and S3D**).

### Male-biased pulmonary IL-1β elevation is associated with neutrophil accumulation

To determine whether the increased neutrophil accumulation observed in male mutant mice was associated with heightened pulmonary inflammatory signaling, *Il1b* expression was quantified in the lungs. Quantitative RT-PCR analysis of whole lung tissue revealed a marked upregulation of *Il1b* transcript levels in *αKl* KO and dKO mice relative to WT and *Rag2* KO controls (**Figure 4A**). Notably, this pulmonary *Il1b* elevation exhibited a pronounced sex bias, with male *αKl* KO and dKO animals displaying higher expression levels than their female counterparts (**Figure 4A**). Combined with the neutrophil accumulation described in Figure 3, these results suggest a potential link between infiltrating neutrophils and increased local IL-1β production. In contrast, the expression levels of other senescence-associated secretory phenotype (SASP)-related and senescence-associated genes assessed in the lungs, including *Tnfa*, *Cdkn2a* (p16), and *Cdkn1a* (p21) (**Figure S4A**), as well as *Il6* (scarcely detectable; data not shown), did not differ significantly between the sexes. These findings indicate that the pulmonary inflammatory phenotype in this model is characterized by a selective increase in *Il1b* expression, but not broad activation of SASP-associated inflammatory or senescence markers.

**Figure 4.**
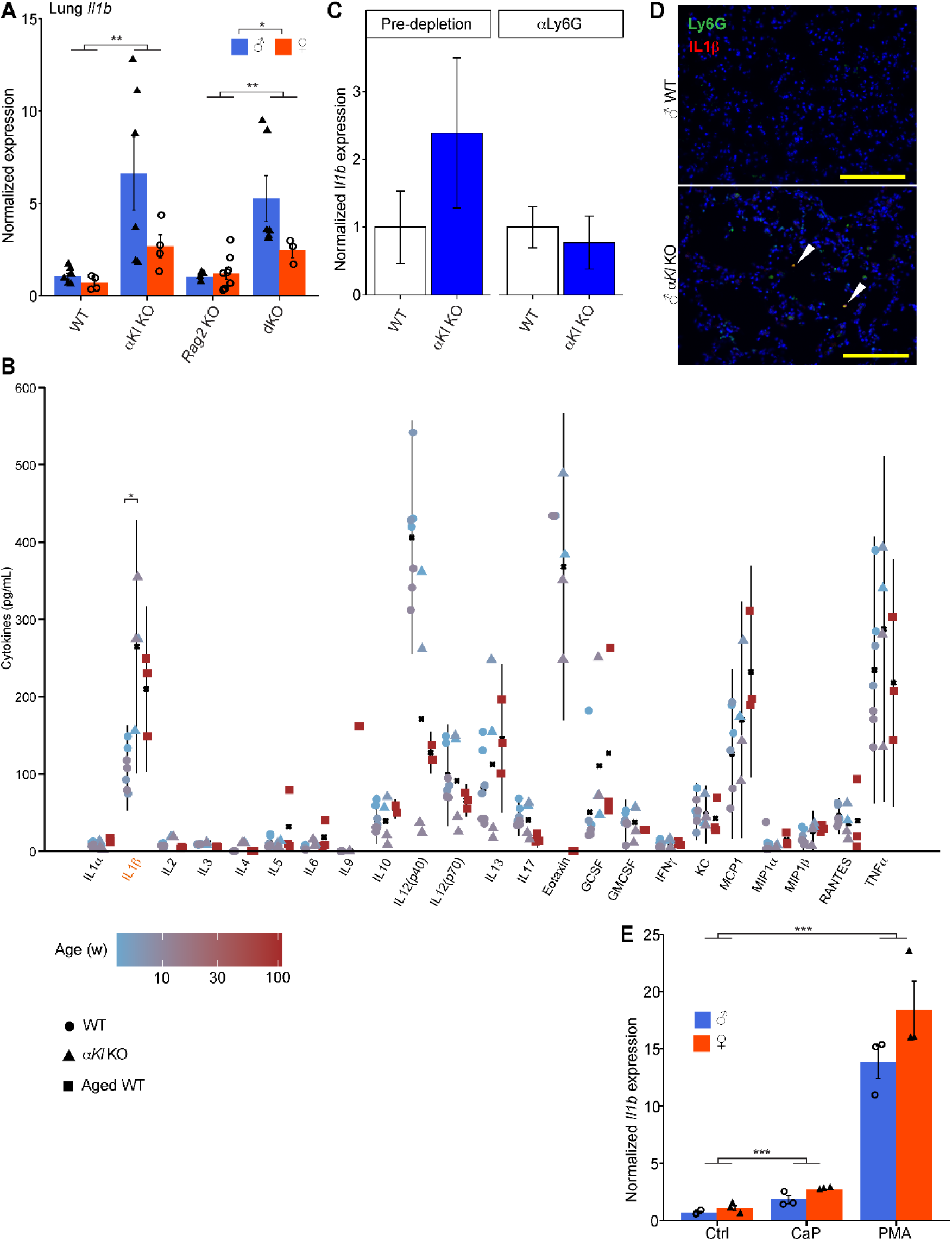
Infiltrating neutrophils drive male-biased pulmonary IL-1β elevation, whereas per-cell cytokine expression remains equivalent across sexes. **(A)** Quantitative RT-PCR analysis of *Il1b* transcript levels in whole lung tissue from male (blue) and female (red) WT, *αKl* KO, *Rag2* KO, and dKO mice (*n* = 3–7 per group). Expression levels are normalized to *Actb*. Data represent mean ± SEM with individual biological replicates shown. Statistical evaluation by two-way ANOVA followed by post-hoc pairwise comparisons with Benjamin-Hochberg adjustment (**: *p* < 0.01 for genotype main effect; *: *p* < 0.05 for sex difference). **(B)** Multiplex serum cytokine profiling across WT (circles), *αKl* KO (triangles), and 2 year-aged WT (squares) mice across a developmental age spectrum (4 to 100 weeks, color gradient). Among 23 quantified cytokines/chemokines, systemic IL-1β (orange label) exhibits selective and significant elevation in KO serum (*: *p* < 0.05, Kruskal-Wallis test followed by Dunn’s test for post-hoc pairwise comparisons with Benjamini-Hochberg adjustment). **(C)** Relative *Il1b* mRNA expression in lung single-cell suspensions prepared from WT (white) and *αKl* KO (blue) male mice before (Pre-depletion, left) and after magnetic depletion of Ly6G^+^ neutrophils (αLy6G, right) (*n* = 3 per group). Data represent mean ± SEM. **(D)** Representative immunofluorescence (IF) images of lung tissue sections from WT (top) and *αKl* KO (bottom) male mice stained for Ly6G (green) and IL-1β (red), with DAPI nuclear counterstain (blue). White arrowheads indicate Ly6G^+^IL-1β^+^ co-localizing neutrophils infiltrating the lung parenchyma of *αKl* KO mice. Scale bars = 100 μm. **(E)** Relative *Il1b* transcript induction in bone marrow-derived neutrophils isolated from male (blue/circles) and female (red/triangles) mice following 4-hour *in vitro* stimulation with control buffer (Ctrl), calcium phosphate (Ca), or phorbol 12-myristate 13-acetate (PMA). Data represent mean ± SEM (*n* = 3 per sex). Statistical evaluation by two-way ANOVA (***: *p* < 0.001 for stimulation main effect; no statistically significant sex difference within treatment groups).

We then assessed whether this localized inflammatory state reflects a broader systemic response across aging by multiplex measurement of cytokines across WT, *αKl* KO, and 2 years-old WT mice. Among the 23 cytokines and chemokines examined, only IL-1β exhibited a selective and significant increase in the circulation of *αKl* KO animals (**Figure 4B**). Notably, this rise was not statistically significant in physiologically aged WT mice, suggesting that elevated IL-1β represents a distinct consequence of *αKl* deficiency rather than a common feature of aging-associated inflammation. Consistent with the tissue-specific nature of this response, *Il1b* expression was not significantly altered by sex in the kidneys of *αKl* KO or dKO mice (**Figure S4B**), indicating that increased IL-1β production is preferentially associated with local pulmonary pathology.

To determine the contribution of neutrophils to pulmonary *Il1b* expression, we isolated single-cell suspensions from the lungs of WT and *αKl* KO male mice before and after magnetic depletion of Ly6G^+^ neutrophils. Consistent with whole-lung measurements, the male *αKl* KO lung single-cell suspension exhibited elevated *Il1b* expression relative to WT control before depletion (**Figure 4C**). However, targeted removal of neutrophils via anti-Ly6G (αLy6G) magnetic depletion completely abolished the elevated *Il1b* signal in *αKl* KO single-cell lung suspensions (**Figure 4C**). These results indicate that infiltrating Ly6G^+^ neutrophils constitute a major source driving the elevated pulmonary *Il1b* expression in mutant mice. Direct visualization of mature IL-1β protein by immunofluorescence staining confirmed these cellular findings: lung tissue sections from male *αKl* KO mice demonstrated marked parenchymal infiltration of Ly6G^+^ neutrophils that robustly co-localized with IL-1β signal (indicated by white arrowheads), whereas WT lungs displayed minimal neutrophil presence and negligible IL-1β expression (**Figure 4D**).

Finally, we tested whether the observed male bias in pulmonary *Il1b* levels stemmed from sexual dimorphism in neutrophil activation capacity or higher neutrophil recruitment to male lungs. Primary bone marrow-derived neutrophils isolated from male and female mice were stimulated *in vitro* with calcium phosphate (CaP) or phorbol 12-myristate 13-acetate (PMA). Both stimuli significantly induced *Il1b* transcript expression, confirming effective neutrophil activation by calcium phosphate precipitates (**Figure 4E**).^34^ However, no statistically significant difference in *Il1b* expression was detected between male and female neutrophils under the tested conditions (**Figure 4E**). These results suggest that both male and female neutrophils have the same inherent ability to increase *Il1b* expression when activated similarly. Together with our neutrophil depletion studies (Figure 4C) and tissue infiltration analyses (Figures 3B and 3C), these data demonstrate that the male-biased elevation of pulmonary *Il1b* observed *in vivo* is driven primarily by increased neutrophil recruitment and accumulation in male lung tissue rather than enhanced cellular reactivity.

### Transcriptomic profiling identifies male-specific dysregulation of extracellular matrix and cell adhesion pathways in bone marrow neutrophils

Having observed the male-biased migratory behaviors of bone marrow-derived neutrophils outside the context of pulmonary inflammation, we performed RNA sequencing on bone marrow neutrophils isolated from male and female *Rag2* KO and dKO mice. Principal component analysis (PCA) demonstrated distinct clustering according to sex (**Figure 5A**). To resolve specific transcriptomic contributions, we mapped overlapping differentially expressed genes (DEGs) across sex (dKO male vs. female; *Rag2* KO male vs. female) and genotype (*Rag2* KO vs. dKO in both sexes) comparisons (**Figures 5B and S5A**‒**C; Tables S1-S4**). The number of differentially expressed genes following *α-Klotho*/*Rag2* deficiency was markedly greater in male than in female neutrophils, suggesting a stronger transcriptional response in males (**Figure S5C**). Among these, male neutrophils showed pronounced transcriptional repression of key structural extracellular matrix (ECM) components, such as *Col11a1*, *Col5a2*, *Sparc*, and *Lgals3bp* (**Figures 5C, S5A, and arrow in Figure 5B**).

**Figure 5.**
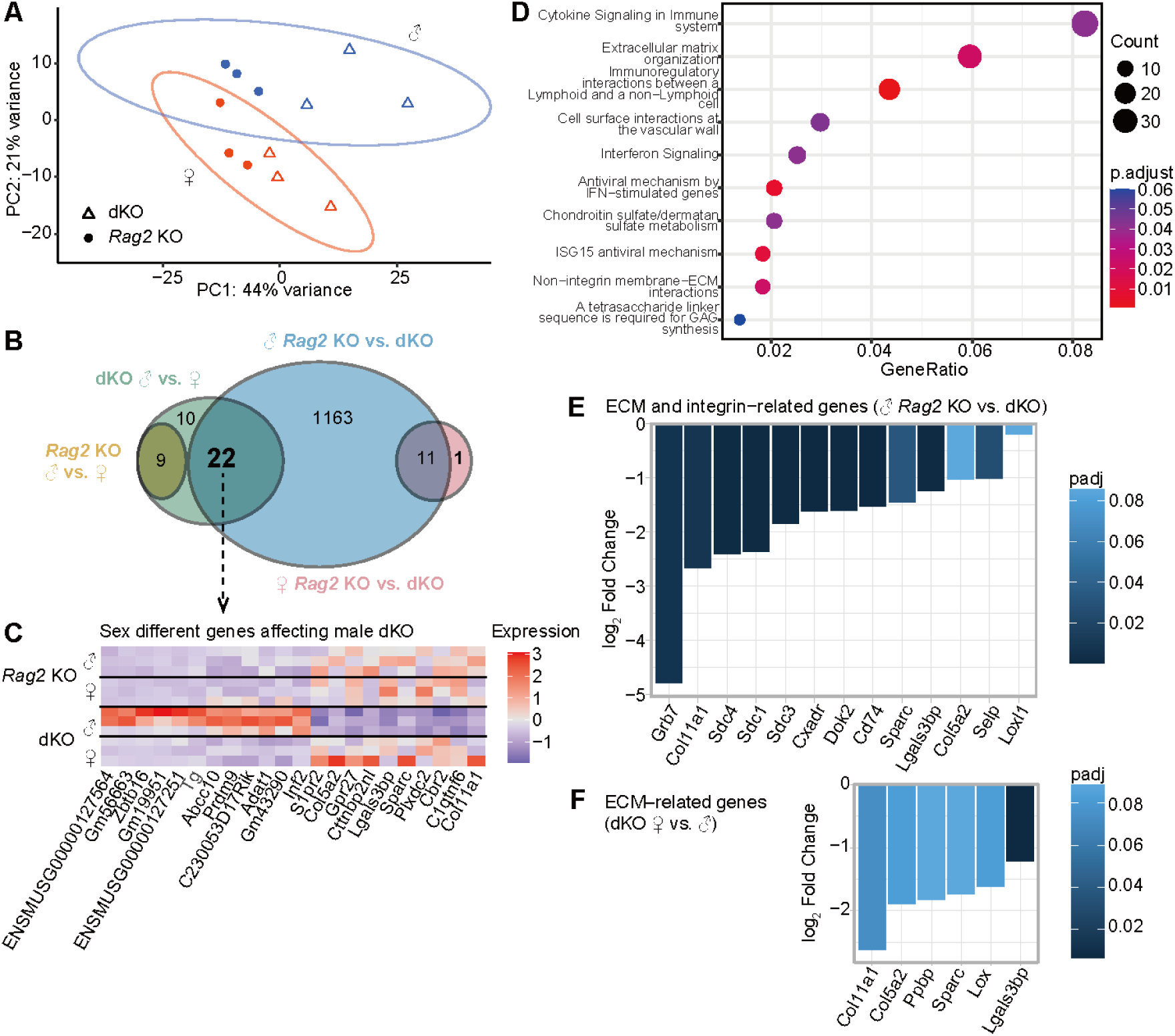
Transcriptomic profiling identifies male-specific dysregulation of extracellular matrix and cell adhesion pathways in bone marrow neutrophils. **(A)** Principal component analysis (PCA) of transcriptomic profiles of bone marrow neutrophils isolated from male (blue) and female (red) Rag2 KO (•) and dKO (△) mice (*n* = 3 mice per group). PC1 and PC2 account for 44% and 21% of the total variance, respectively. Ellipses depict the 95% confidence intervals. **(B)** Venn diagram showing the overlap of differentially expressed genes (DEGs) across sex comparisons (dKO male vs. female [green]; *Rag2* KO male vs. female [yellow]) and genotype comparisons (male *Rag2* KO vs. dKO [blue]; female *Rag2* KO vs. dKO [pink]). A core set of 22 sex-differentiated DEGs shared across male *Rag2* KO vs. dKO as well as dKO male vs. female comparisons is highlighted (dashed arrow). **(C)** Heatmap displaying the relative expression (z-score) of the 22 sex-differentiated genes across *Rag2* KO and dKO male vs. female neutrophils. **(D)** Pathway enrichment analysis (Reactome/GO dot plot) of altered biological processes in male neutrophils. The circle size corresponds to the gene count within each pathway, and the fill color indicates the adjusted *p*-value (p.adjust). **(E)** log_2_ fold change of ECM- and integrin-related genes comparing male *Rag2* KO versus male dKO neutrophils. Negative values denote significant downregulation in the dKO state. The bar fill color represents the adjusted *p*-value (padj). **(F)** log_2_ fold change of key ECM-associated genes comparing female versus male dKO neutrophils (*Col11a1*, *Col5a2*, *Ppbp*, *Sparc*, *Lox*, and *Lgals3bp*). Negative values indicate lower relative expression of matrix-anchoring genes in male dKO neutrophils. The bar fill color represents the adjusted *p*-value (padj).

Pathway enrichment analysis was performed to gain insights into the biological processes represented by these transcriptional changes (**Figure 5D**). Among the pathways enriched in male neutrophils, *extracellular matrix organization* and *cell surface interactions at the vascular wall* were remarkable (**Figure 5D**). Direct assessment of effect sizes (log_2_ fold change) confirmed significantly decreased expression of core ECM-associated genes (*Lgals3bp*, *Sparc*, and *Col11a1*) as well as integrin-related genes (*Grb7*, *Sdc1*, *Sdc3*, *Sdc4*, *Cxadr*, *Dok2*, *Cd74*, and *Selp)*, when comparing male *Rag2* KO and dKO neutrophils (**Figure 5E**). Moreover, comparison of male and female dKO neutrophils shared the modulation of ECM-related genes (**Figure 5F**), indicating that both sex and *αKl* deficiency influenced pathways associated with extracellular matrix interactions.

Taken together, these transcriptomic analyses suggest that neutrophils undergo preferential remodeling of pathways regulating cell adhesion, which is associated with sex as well as *αKl* deficiency, consistent with altered migratory behavior.

### Sex-biased alternative splicing enriches Toll-like receptor signaling pathways, with hormones modulating integrin isoform usage

Given the absence of *Itgam* (coding CD11b) among the sex-dependent DEGs identified in *Rag2* KO or dKO neutrophils, we investigated whether alternative splicing (AS) contributes to the general mechanism underlying sexual dimorphism in neutrophil biology. Re-analysis of public RNA-sequencing datasets from WT neutrophils^25^ revealed widespread sex-biased AS events across multiple event categories, including skipped exons (SE), mutually exclusive exons (MXE), alternative 3’ and 5’ splice sites (A3SS, A5SS), and alternative first and last exons (AFE, ALE) (**Figure 6A and S6A; Table S5**). Reactome pathway enrichment analysis of these differentially spliced transcripts in WT neutrophils demonstrated marked enrichment in pattern recognition pathways, specifically the *Toll-Like Receptor 4 (TLR4) cascade*, *TRIF(TICAM1)-mediated TLR4 signaling*, and the *MyD88-independent TLR4 cascade* (**Figure 6B**). Similar enrichment of TLR cascades, as well as mRNA processing pathways, was observed in 20-month-old WT neutrophils (**Figures S6C–D; Table S6**). These results suggest that sex-biased alternative splicing may preferentially affect the components of innate immune sensing pathways and potentially contribute to functional differences in neutrophil activation.

**Figure 6.**
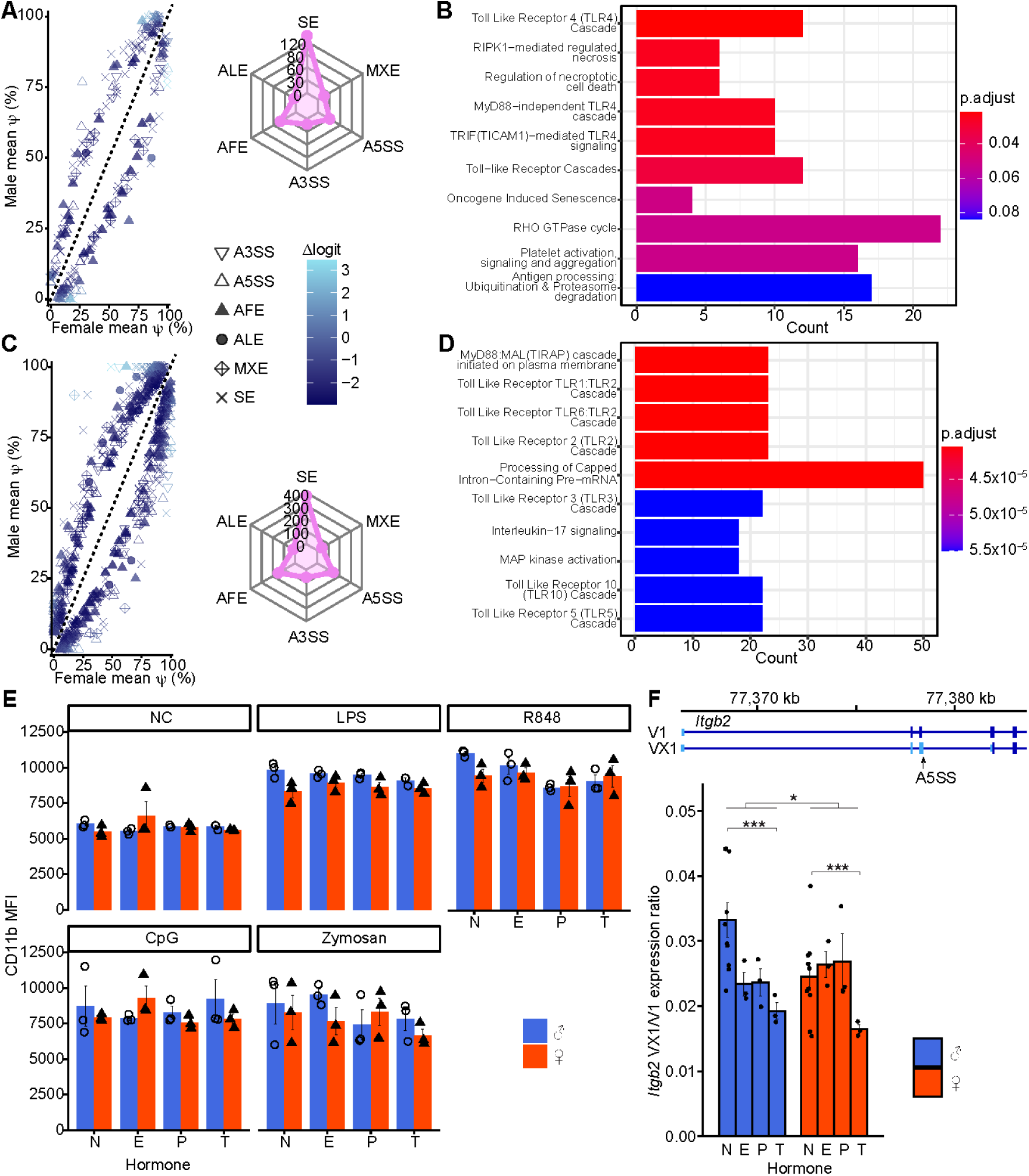
Sex-biased alternative splicing enriches Toll-like receptor signaling pathways, with hormones modulating *Itgb2* isoform usage. **(A)** Distribution of sex-biased alternative splicing (AS) events in WT bone marrow neutrophils re-analyzed from public RNA-seq datasets, plotting male mean Percent Spliced In (ψ (%)) against female mean ψ (%). The categorized splicing event types include alternative 3’ splice sites (A3SS), alternative 5’ splice sites (A5SS), alternative first exons (AFE), alternative last exons (ALE), mutually exclusive exons (MXE), and skipped exons (SE). Point shading indicates the magnitude of the logit inclusion shift (|Δlogit|). The radar chart on the right shows the number of events per alternative splicing modality. **(B)** Reactome pathway enrichment analysis of differentially spliced transcripts in male versus female WT neutrophils. The bar length indicates the gene count, and the fill color corresponds to the adjusted *p*-value (p.adjust). Prominent enrichment was observed in the *Toll-Like Receptor 4 (TLR4) cascade*, *TRIF(TICAM1)-mediated TLR4 signaling*, and *MyD88-independent TLR4 cascade*. **(C)** Alternative splicing landscape in bone marrow neutrophils from *Rag2* KO mice (own RNA-seq dataset), plotting male mean ψ (%) versus female mean ψ (%), with events colored by change in log-odds (Δlogit). The radar chart on the right shows the event count per alternative splicing modality. **(D)** Reactome/GO pathway enrichment analysis of spliced gene sets in *Rag2* KO neutrophils showing significant enrichment across TLR family networks (TLR1/2, TLR2, TLR6/2, TLR3, TLR5, TLR10 cascades), IL-17 signaling, and MAP kinase activation (p.adjust indicated by color gradient). **(E)** Surface CD11b expression (mean fluorescence intensity, MFI) in male (blue) and female (red) bone marrow neutrophils following ex vivo stimulation with vehicle control (NC) or distinct TLR agonists—LPS (TLR4), R848 (TLR7/8), CpG (TLR9), or zymosan (TLR2)—in the presence or absence of sex hormones: None (N), 17β-estradiol (E), progesterone (P), or testosterone (T) (*n* = 3 per group). Data are presented as mean ± SEM with individual biological replicates shown. **(F)** Schematic representation of the integrin β2 (*Itgb2*) genomic locus showing alternative transcripts V1 and VX1 (top). Light blue indicates the 5’ UTR. Quantitative RT-PCR assessment of the *Itgb2* VX1/V1 expression ratio (bottom) in male (blue) and female (red) bone marrow neutrophils treated with sex hormones (N, E, P, and T) (*n* = 3–10 per group). Data are presented as mean ± SEM, with individual data points shown. Statistical evaluation by two-way ANOVA followed by post-hoc pairwise comparisons with Benjamin-Hochberg adjustment (***: *p* < 0.001 for hormonal main effect; *: *p* < 0.05 for sex differences).

To confirm these post-transcriptional patterns in our experimental model, we mapped the alternative splicing landscape of bone marrow neutrophils from *Rag2* KO mice (**Figure 6C and S6B; Table S7**). Similar to the analysis of WT neutrophils, sex-associated splicing differences were observed across a wide range of splice event types (**Figure 6C**). Pathway enrichment analysis of spliced gene sets in *Rag2* KO neutrophils mirrored the public dataset, demonstrating pronounced enrichment across broader TLR family networks, including TLR1/2, TLR2, TLR6/2, TLR3, TLR5, and TLR10 cascades (**Figure 6D**). Enrichment of mRNA processing and splicing components was also observed in female versus male neutrophils of dKO mice (**Figures S6E–F; Table S8**). Collectively, these analyses support the hypothesis that alternative splicing represents an additional layer of regulation influencing inflammatory signaling networks in male and female neutrophils.

Then, do sex and sex hormones regulate CD11b expression or finely tune the alternative splicing of TLR pathway-related genes? We measured the surface expression of CD11b following *ex vivo* stimulation with distinct TLR agonists in the presence or absence of 17β-estradiol (E), progesterone (P), or testosterone (T). As expected, stimulation with TLR ligands increased CD11b surface expression relative to unstimulated controls; however, the magnitude of this response varied according to both sex and hormone treatment (**Figure 6E**).

Because cell surface mobilization and integrin function rely on specific structural subunit variants, we investigated whether sex hormones directly regulate splice site selection in integrin β2 (*Itgb2*), the gene encoding the integrin subunit that forms the Mac-1 integrin complex together with CD11b, which is also a downstream molecule in TLR signaling pathways. Quantitative RT-qPCR assessment of the expression ratio of *Itgb2* alternative splicing isoforms (VX1/V1) revealed sex-specific baseline differences (**Figure 6F**). Furthermore, testosterone treatment altered isoform usage patterns, suggesting that sex hormones may influence the post-transcriptional regulation of integrin expression. Although the functional consequences of these isoform shifts remain to be established, these data raise the possibility that sex-dependent alternative splicing contributes to the regulation of neutrophil adhesion and activation pathways.

Collectively, these findings indicate that sex-biased alternative splicing is a prominent feature of bone marrow neutrophils and preferentially affects genes associated with innate immune signaling networks. The enrichment of TLR-related pathways, together with hormone-responsive modulation of *Itgb2* isoform usage and male-biased surface CD11b expression, suggests a potential link between sex-dependent RNA processing and the regulation of neutrophil migratory behavior.

## Discussion

The prevailing view of immunological sex differences has largely focused on divergent responses to infection, autoimmunity, and hormonal stimulation after immune activation. Previous studies on sex bias of neutrophils have largely focused on stimulus-induced activation signals.^25,35^ In contrast, our findings suggest that sexual dimorphism may already be encoded within the resting neutrophil compartment. In this study, we identified a previously underappreciated male-biased neutrophil state characterized by elevated basal surface expression of integrin CD11b and associated transcriptional and post-transcriptional programs, independently of adaptive lymphocytes or overt inflammatory stimulation. These findings support a model in which inflammatory susceptibility is at least partially encoded within the resting innate immune compartment before pathological activation occurs, extending recent evidence that neutrophils exhibit sex-dependent developmental and functional heterogeneity.^36^ CD11b is widely regarded as a marker of neutrophil activation; however, Mac-1 signaling also controls adhesion, intravascular crawling, endothelial interactions, and tissue infiltration.^11,16^ Our findings suggest that physiological variations in baseline CD11b abundance, even in the absence of overt inflammatory stimulation, may influence the threshold for tissue recruitment of neutrophils. This interpretation differs from the classical view that integrin upregulation is merely a consequence of activation and instead suggests that integrin availability itself may define distinct neutrophil states. Additionally, a recent study showing different responses to chemorepellents between male and female human neutrophils^37^ parallels our interpretation that adhesion and migration behaviors are distinctly regulated between sexes.

An intriguing aspect of our study was the organ specificity of the observed phenotype. According to publicly available human single-cell expression data, lung epithelial cells particularly possess substantial expression of *ICAM1* (binding partner for Mac-1 integrin), indicating the highest frequencies of integrin-mediated neutrophil attachment and subsequent infiltration.^38^ Consequently, relatively modest alterations in neutrophil adhesive programs may disproportionately affect pulmonary leukocyte retention and tissue infiltration. Recent work has emphasized that biological sex strongly influences lung immunity and inflammatory responses, yet the cellular and molecular mechanisms remain incompletely defined.^39^ Our findings raise the possibility that a sex-dependent adhesive neutrophil state contributes to organ-specific inflammatory vulnerability by selectively enhancing pulmonary retention and recruitment.

The identification of altered neutrophil phenotypes in physiologically aged wild-type mice^25^ suggests that this male-biased inflammatory program is not unique to *α-Klotho* deficiency. Aging is increasingly recognized as a process accompanied by chronic low-grade inflammation and remodeling of myeloid cell populations.^10,40^ Several studies have described the accumulation of specialized neutrophil states during aging, including senescence-associated and chronologically aged neutrophil populations exhibiting altered migratory and inflammatory properties.^10,41^ Our results extend these observations by suggesting that biological sex may overlay an additional molecular regulatory network of neutrophil heterogeneity interacting with aging-associated immune remodeling. Importantly, however, the progeric phenotype observed in *α-Klotho* - deficient mice cannot be explained solely by physiological ageing, indicating that accelerated mineral metabolism-associated disease and natural aging remain biologically distinct conditions despite sharing certain immunological features.

Unexpectedly, conventional differential gene expression analysis did not identify obvious pathways directly explaining the increased CD11b surface abundance. Instead, reduced expression of extracellular matrix- and adhesion-associated genes was noted only in the *αKl*-deficient disease setting. Although direct measurements of neutrophil retention were not performed, this finding is consistent with the evidence that neutrophil trafficking is regulated not only by chemotactic cues but also by interactions with stromal niches and extracellular matrix architecture. Alterations in these pathways may influence the bone marrow niche for neutrophil mobilization,^42^ thereby contributing to sex-dependent migratory behaviors. Future studies directly examining neutrophil migration *in vivo* are necessary to test this possibility.

The most striking observation was that sex-dependent differences were more prominent at the level of alternative splicing than that of conventional gene expression. Alternative splicing has emerged as an important mechanism for generating functional diversity within immune cells, enabling the fine regulation of receptor signaling, inflammatory responses, and cellular activation thresholds.^20,22^ Our findings further raise the possibility that sex-dependent RNA processing contributes to the establishment of distinct neutrophil activation states, consistent with recent proteomic evidence suggesting enhanced RNA-processing and signaling programs in unstimulated male neutrophils.^43^ As for an interpretation of our results, sequence analysis using SignalP 6.0^44^ revealed a decisive functional distinction: whereas the canonical Itgb2 V1 isoform harbors a conserved *N*-terminal signal peptide essential for co-translational translocation into the endoplasmic reticulum and subsequent cell surface trafficking, the VX1 isoform lacks an entire signal peptide (**Table S9**). These predictions may contribute to linking alternative splicing to altered integrin availability, protein trafficking, and cell-surface expression, potentially influencing distinct neutrophil states.

Acute exposure to sex hormones was insufficient to fully recapitulate the male-associated splicing pattern, suggesting that sex-biased neutrophil states are unlikely to be explained solely by short-term hormonal signaling. Instead, long-term developmental, epigenetic, or hormonally programmed regulatory mechanisms may establish stable sex-specific RNA-processing programs.^45^ Similar conclusions have emerged from recent studies indicating that sex-linked immune programs can persist beyond immediate endocrine regulation and may involve developmental, epigenetic, or chromatin-based mechanisms. Taken together, our findings suggest that sex-dependent neutrophil states under resting conditions are established through long-term programming processes that subsequently influence inflammatory responsiveness and disease susceptibility throughout an individual’s life.

### Limitations of the study

This study has several limitations. First, the human analyses were derived from inflammatory disease cohorts and lacked adequately represented healthy male controls, limiting the conclusions regarding physiological human neutrophil dimorphism (**Figure S2C**). Second, although alternative splicing analyses identified sex-biased *Itgb2* isoforms and TLR-associated programs, the causal contribution of specific splice isoforms to CD11b surface abundance remains to be established. Third, CD11b blockade was evaluated *in vitro*, and future studies are required to determine whether the manipulation of neutrophil trafficking can modify pulmonary pathology *in vivo*. Finally, direct comparisons between murine bone marrow neutrophils and human circulating granulocytes should be interpreted cautiously because of the potential compartment-specific differences.

## Conclusion

In summary, our findings support a model in which biological sex influences the resting neutrophil state associated with differential integrin availability and RNA-processing programs. The identification of a male-biased CD11b^hi^ neutrophil state provides a potential explanation for the altered tissue recruitment and inflammatory susceptibility. Rather than surveying immunity after overt activation, this study provides a framework for understanding sex-specific differences in disease vulnerability preconfigured within resting neutrophils across both pathological and physiological contexts.

## Resource availability

### Lead Contact

Requests for further information and resources should be directed to and will be fulfilled by the lead contact, Mayumi Mori.

### Materials Availability

Resources are available upon request to the lead contact. Supplementary figures are found in a separated file.

### Data and Code Availability

The RNA sequencing data generated in this study have been deposited in the Gene Expression Omnibus (GEO) under accession number GSE353159, within NCBI BioProject database (accession number: PRJNA1554482). The data remain private during the peer review and may be accessed using the reviewer token provided by GEO. Code for computing log-odds ratios and fitting curves is available upon request. Publicly available sequencing data were obtained from the NCBI BioProject database (accession number: PRJNA630663). The human public flow cytometry data re-analyzed in this study were obtained from ImmPort (https://www.immport.org) under study accession SDY9, “ADVN Biomarker Registry: Neutrophil Substudy,”

## Acknowledgments

We express our gratitude to Prof. Shigeo Koyasu, National Institutes for Quantum Science and Technology (QST), for providing *Rag2* KO mice through the Center for Animal Resources and Development (CARD), Kumamoto University, and to Dr. Quan Wu, Kyoto University, for generously providing aged wild-type mice (supported by Japan Science and Technology Agency (JST), Precursory Research for Embryonic Science and Technology (PRESTO) / Sakigake JPMJPR2382) as well as sharing lab equipment. We also acknowledge the Center for Anatomical Studies, Kyoto University Graduate School of Medicine, for technical assistance with histological and immunohistochemical analyses. RNA sequencing library preparation and sequencing were supported by the Single-cell Genome Information Analysis Core (SignAC) at the Institute for Advanced Study of Human Biology (WPI-ASHBi), Kyoto University. Flow cytometry acquisition and analysis were performed using equipment at the Medical Research Support Center (iSAL), Kyoto University Graduate School of Medicine. We thank Ms. Miwa Washida for technical assistance with mouse husbandry and survival assessment, Prof. Yoko Hamazaki, Center for iPS Cell Research and Application (CiRA), Kyoto University, for technical support and assistance with Bio-Plex cytokine profiling, and Dr. Yuki Kanesaka for technical assistance with quantitative RT-qPCR. In addition, we extend our sincere thanks to Prof. Fumio Matzuzaki, Kyoto University, and Dr. Quan Wu for valuable scientific discussions and advice throughout this study. This work was supported by Grant-in-Aid for Challenging Research (Exploratory) / 25K22776 to M.M. This work was conducted at the Endowed Chair of Department of Aging Science and Medicine, Kyoto University, under the direction of Specially Appointed Professor Y-I.N. This department is financially supported by generous donations from Meiji Holdings Co., Ltd., Gakken Holdings Co., Ltd., and NTT, Inc. The funders had no role in the study design, data collection and analysis, decision to publish, or preparation of the manuscript.

## Author contributions

M.M. conceptualized the study, designed and performed most experiments, analyzed data, secured funding, and wrote the manuscript. C.A. contributed to conceptual research design, performed experiments, and analyzed data for cytokine profiling and longitudinal physical tracking. S.S. executed the primary bioinformatic alternative splicing pipeline under the supervision of T.Y.. T.T. conducted RNA-sequencing experiments and transcriptomic data processing. S.M. provided single-knockout survival cohort data under the supervision of H.M.. Y-I.N. acquired funding, supervised the overall research project, and edited the manuscript. All authors reviewed and approved the final manuscript.

## Declaration of generative AI and AI-assisted technologies in the manuscript preparation process

During the preparation of this work, the authors used Paperpal and generative AI tools (Microsoft 365 Copilot and Google Gemini (ver.3.5 Thinking and ver.3.6 Thinking)) in order to assist in manuscript organization and text refinement. After using this tool/service, the authors reviewed and edited the content as needed and take full responsibility for the content of the published article.

## Methods

Sources and identifiers for biological materials, reagents, and software are found in **Table S10.**

### Mice and Animal Models

*α-Klotho* KO (*αKl* KO) mice on a C57BL/6J background were previously generated.^26,27^ To generate lymphocyte-deficient double-knockout (dKO) animals, heterozygous *αKl^+/-^* mice were crossed with *Rag2* KO (*Rag2^-/-^*) mice. *Rag2^+/-^αKl^+/-^* offspring were intercrossed to yield wild-type (WT), *αKl* KO, *Rag2* KO, and dKO (*αKl*^-/-^*Rag2*^-/-^) littermates. All mice were housed in a specific pathogen-free (SPF) facility under a standard 12-hour light/dark cycle at 22–24°C and 40–60% humidity, with free access to standard chow and water. Animal experiment protocols were approved by the Institutional Animal Care and Use Committee.

### Survival Analysis and Longitudinal Physical Tracking

Mice were monitored daily for health status and survival. For survival analysis, the endpoint criteria included the loss of righting reflex, severe lethargy, or moribund state. Median survival was evaluated using Kaplan–Meier curves, and statistical significance between sexes and genotypes was assessed using log-rank (χ^2^) and Gehan–Wilcoxon tests. Body weight, serum phosphorus, and serum calcium levels were tracked longitudinally from 6 to 12 weeks of age in male and female *Rag2* KO and dKO animals.

### Histology, Pulmonary Pathology, and Tissue Calcification Assay

Lungs were harvested following systemic vascular perfusion with ice-cold PBS. Tissues were fixed in 10% neutral formalin for 24 hours at 4°C, embedded in paraffin, and sectioned at a thickness of 4 μm. Hematoxylin and eosin (H&E) staining was performed according to standard procedures to evaluate pulmonary emphysema and parenchymal structural integrity.

Pulmonary calcium deposition was quantified using an Alizarin Red S colorimetric assay. Lung tissues were homogenized in 4% paraformaldehyde (PFA) in PBS, cleared by centrifugation at 13,000 × *g* for 5 minutes at 4°C, and incubated with 2% w/v Alizarin Red S solution (40 mM, pH 4.2). After extraction with 10% acetic acid for 30 minutes at room temperature + 10 minutes at 85°C, absorbance was measured at 450 nm using a microplate reader. Tissue calcium concentration was calculated against a standard curve of 0.0313 ∼ 2 mM Alizarin Red S solution and normalized to tissue weight (μmol/mg tissue).

### Serum Biochemistry and Cytokine Profiling

Blood was collected via cardiac puncture, allowed to clot at room temperature for 30 minutes, and centrifuged at 2,000 × *g* for 10 minutes at 4°C to isolate the serum. Samples were sent to SRL for outsourced analysis of the serum concentrations of 1,25-(OH)_2_D_3_ (vitamin D). Serum inorganic phosphorus and calcium levels were quantified using colorimetric assay kits. Multiplex profiling of 23 systemic cytokines and chemokines across age groups (4–100 weeks) was conducted using a Bio-Plex Pro Mouse Cytokine 23-plex Panel on a Bio-Plex 200 System.

### Primary Cell Isolation and Flow Cytometry

Bone marrow cells were harvested by flushing the femurs and tibias with ice-cold PBS. Lung and kidney tissues were minced and digested with Liberase TM (0.13 mg/mL) and DNase I (10 μg/mL) at 37°C for 45 minutes, followed by mechanical dissociation through a 70-μm cell strainer. Red blood cells were lysed using ACK Lysis Buffer.

Single-cell suspensions were blocked with anti-CD16/32 (Fc block) for 10 minutes on ice and incubated with fluorophore-conjugated antibodies for 30 minutes at 4°C in the dark. The following cell populations within viability dye-negative gating were defined:

#### Monocytes or neutrophil progenitors and mature neutrophils

Ly6C^hi^CD115^+^ cells were gated as monocyte progenitors. Ly6C^+^CD115^-^CD11b^+^Ly6G^+^ neutrophil progenitors were further gated on CD101^+^ mature neutrophils.

#### Lung leukocyte subsets

CD45^+^ total leukocytes, Ly6G^+^CD11b^+^CD11c^-^neutrophils, CD11c^+^Siglec-F^+^ alveolar macrophages, CD11c^-^Siglec-F^+^ eosinophils, CD11b^+^F4/80^+^ interstitial macrophages, CD4^+^ T cells, and CD8^+^ T cells.

#### Renal immune subsets

Total renal CD45^+^ leukocytes and Ly6G^+^CD11b^+^ neutrophils.

CD11b mean fluorescence intensity (MFI) was evaluated in bone marrow neutrophils. Flow cytometric acquisition was performed using a BD LSRFortessa, and the data were analyzed using FlowJo software.

### In Vitro Neutrophil Transmigration Assay

Endothelial transmigration assays were performed using Transwell inserts (24-well plate, 3.0 μm pore size).^32,33^ Mouse brain microvascular endothelial cells (bEnd.3) were seeded onto the upper chambers and cultured to confluence. Prior to the assay, bEnd.3 monolayers were treated with TNFα (20 ng/mL) for 4 hours, after which the medium was changed to serum-free RPMI-1640. Primary bone marrow neutrophils were isolated from male and female WT mice using the EasySep mouse neutrophil enrichment kit. Purity was assessed by Ly6G^+^CD11b^+^ together with CD115 staining on flow cytometry to confirm minimal contamination of monocytes. Isolated cells were resuspended in serum-free RPMI-1640 medium and pre-treated with either anti-CD11b neutralizing antibody (5 μg/mL) or an isotype control IgG for 15 minutes on ice. Neutrophils (2 × 10^5^ cells) were added to the upper chamber, and recombinant mouse CXCL2 (50 ng/mL) was added to the lower chamber as a chemoattractant. After 45 minutes of incubation at 37°C, the number of migrated Ly6G^+^ neutrophils in the bottom well was quantified by flow cytometry and normalized to gravity-dependent baseline transmigration without chemoattractant.

### Immunohistochemistry (IHC) and Immunofluorescence (IF)

Immunohistochemical staining was performed using the conventional avidin-biotin-complex (ABC) method.^46^ Tissue sections were deparaffinized and subjected to antigen retrieval using a microwave oven with citrate buffer for 15 minutes. Endogenous peroxidase activity was blocked by incubating the glass slides in 0.3% hydrogen peroxide in methyl alcohol for 30 minutes. Following blocking, the slides were washed in PBS six times for 5 minutes each and subsequently mounted with 1% normal serum in PBS for 30 minutes. The primary antibody against Ly-6G (Proteintech: 65078-1-Ig, rat IgG, diluted 1:200) was applied overnight at 4°C. After washing in PBS (six times, 5 minutes each), a biotinylated secondary antibody diluted to 1:300 in PBS was applied for 40 minutes. The slides were then washed in PBS (six times, 5 minutes each) and incubated with the avidin-biotin-peroxidase complex (ABC-Elite, Vector Laboratories, Burlingame, CA) diluted 1:100 in 1% bovine serum albumin (BSA) in PBS for 50 minutes. After a final wash in PBS (six times, 5 minutes each), the color reaction was developed using 3,3’-diaminobenzidine (DAB), and the nuclei were counterstained with hematoxylin. All incubation and washing procedures were carried out at room temperature unless otherwise specified.

Alternatively, for immunofluorescence (IF), primary antibodies against Ly-6G (Proteintech: 65078-1-Ig, rat IgG, diluted 1:200) and IL-1β (Invitrogen: MA5-47038, rabbit IgG, diluted 1:500) were applied simultaneously overnight at 4°C. After washing in PBS (six times, 5 minutes each), fluorescent-conjugated secondary antibodies were applied for 40 minutes at room temperature: Alexa Fluor 488-conjugated anti-rat secondary antibody (1:500 dilution) for Ly-6G detection and Alexa Fluor 594-conjugated anti-rabbit secondary antibody (1:500 dilution) for IL-1β detection. The slides were then washed in PBS (six times, 5 minutes each). Nuclear counterstaining was performed using DAPI (4’,6-diamidino-2-phenylindole). After a final wash in PBS, slides were mounted with an appropriate anti-fade mounting medium and examined under a fluorescence microscope. All incubation and washing procedures were carried out at room temperature unless otherwise specified. Fluorescence images were acquired using KEYENCE BZ-X800.

### Neutrophil Magnetic Depletion and Ex Vivo Cell Stimulation

Single-cell suspensions prepared from male WT and *αKl* KO lungs were incubated with anti-Ly6G antibody, then neutrophils were depleted by streptavidin-coated magnetic beads using an EasySep magnetic stand. Unfractionated and neutrophil-depleted single-cell suspensions were harvested for total RNA extraction.

For functional stimulation, primary bone marrow neutrophils were isolated, resuspended in PBS/G-CM (PBS + 0.2% _D_-glucose + 0.9 mM CaCl_2_ + 0.5 mM MgCl_2_),^34^ and stimulated with calcium phosphate precipitates (CaP, 10 mM) or PMA (100 nM). For TLR stimulation, neutrophils were resuspended in RPMI-1640 with charcoal-stripped fetal bovine serum and cultured in the presence or absence of 17β-estradiol (10 pM), progesterone (1 nM), or testosterone (2 μM). After 20 hours of culture, neutrophils were incubated with TLR agonists—LPS (10 ng/mL, TLR4), R848 (125 nM, TLR7/8), CpG (2 μM, TLR9), or Zymosan (1 μg/mL, TLR2)—for 30‒45 minutes at 37°C.

### RNA Isolation and Quantitative RT-PCR Analysis

Total RNA was extracted from homogenized lung/kidney tissues using TRIzol Reagent and Direct-zol RNA Miniprep (Zymo). Bone marrow neutrophils were preserved in RNAlater immediately upon isolation, then total RNA was extracted by RNeasy Plus Mini kit. cDNA was synthesized using the High-Capacity cDNA Reverse Transcription Kit or High-Capacity RNA-to-cDNA Kit. Quantitative real-time PCR was performed using a StepOnePlus system with SYBR Green Master Mix. Relative transcript expression of *Il1b*, *Tnfa*, *Cdkn2a*, and *Cdkn1a* was calculated via the 2^-ΔΔ^ ^CT^ method and normalized to *Actb*. Integrin β2 (*Itgb2*) alternative splicing isoform ratios (VX1/V1) were quantified using isoform-specific reverse primers and a common forward primer.

### RNA Sequencing, Differential Expression, and Alternative Splicing (AS)

Total RNA was isolated from bone marrow neutrophils of male and female *Rag2* KO and dKO mice (*n* = 3 mice per group). Ribosomal RNA depletion was performed using 100 ng of total RNA, followed by library preparation using the Illumina Stranded Total RNA Prep and ligation with Ribo-Zero Plus kit (cat# 20040525). Sequencing was performed on an Illumina NovaSeq 6000 platform in 151 bp paired-end mode. Adapter sequences and low-quality bases were trimmed from the raw reads using cutadapt v4.6. The trimmed reads were mapped to mouse reference genome sequences (mm39) using STAR v2.7.11a with the GENCODE (GENCODE vM33/Ensembl110 annotations) gtf file. Raw counts were calculated using htseq-count v2.0.5 with the GENCODE gtf file. Differential gene expression analysis was conducted using DESeq2 v1.42.0. Differentially expressed genes (DEGs) were defined by padj < 0.1. Pathway enrichment analysis (Reactome and Gene Ontology) was performed using the clusterProfiler package (v4.8.2). Other packages used were ReactomePA v1.44.0 and fgsea v1.26.0.

Alternative splicing events, including skipped exons (SE), mutually exclusive exons (MXE), alternative 3’/5’ splice sites (A3SS/A5SS), and alternative first/last exons (AFE/ALE), were quantified using percent spliced-in (PSI) values, as previously described.^47^ Additionally, based on Percent Spliced In (ψ = PSI) and |Δψ| (%), log-odds ratios (Δlogit) were calculated as ln(odds_male_) – ln(odds_female_) = ln(odds_male_/odds_female_), where odds = ψ / (1 - ψ). Significantly differentially spliced sites were defined as |Δψ| > 0.05, total junction reads > 20, and Δlogit > ln2 (= 0.693). According to Equations 1–2 of a reported scaling law for splicing,^48^ exon inclusion is modeled as competing kinetic processes governed by splice-site rates k and k_0_. When a mutation or condition change alters relative splicing kinetics by a factor of ð = Δ ln(*k*/*k*_0_), the new ψ as a function of the baseline control ψ_0_ is:

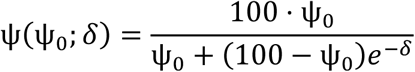

This equation was used for the fitting curves of |Δψ| (%) versus ψ (%) plots. Public RNA-seq datasets of young and 20-month-old wild-type mouse bone marrow neutrophils^25^ were re-analyzed using the same bioinformatic pipeline.

### Human Patient Cohort Bioinformatic Re-analysis

Public flow cytometric and transcriptomic datasets derived from circulating granulocytes and non-granulocytes of healthy human controls (HC) and patients with atopic dermatitis (AD) or AD complicated by eczema herpeticum (ADEH) were re-analyzed. Granulocytes were gated on the CD16^+^SSC^hi^ population and non-granulocytes including lymphocytes and monocytes as the second largest population were gated on the CD16^lo^SSC^lo^ population. Surface CD11b MFI values were extracted, stratified by biological sex, and statistically compared using two-way ANOVA with Benjamini-Hochberg post-hoc correction.

### Statistical Analysis

Statistical analysis was performed using R. Continuous data are presented as mean ± SEM or median with interquartile range. Survival distributions were evaluated using log-rank and Gehan–Wilcoxon tests. Group comparisons involving sex, genotype, or treatment factors were analyzed by two-way ANOVA using ARTool package (v0.11.2), unless specified. Correlations were calculated using Spearman’s rank correlation (ρ) and linear regression (β_age_). Statistical significance was defined as *p* < 0.05 (*: *p* < 0.05, **: *p* < 0.01, ***: *p* < 0.001).

