## Supplementary figures for "Alternative splicing links sex to neutrophil inflammatory vulnerability"

[5]Lead contact

Correspondence to:

Mayumi Mori,

**Supplementary Table Titles and Figures**

**TableS1_Rag2_FvsM_DEG_FDR0_1**

**TableS2_dKO_FvsM_DEG_FDR0_1**

**TableS3_Mrag2_vs_dko_DEG_FDR0_1**

**TableS4_Frag2_vs_dKO_DEG_FDR0_1**

**TableS5_AS_FM5m_LogitFiltered**

**TableS6_AS_FM20m_LogitFiltered**

**TableS7_AS_Rag2_LogitFiltered**

**TableS8_AS_dKO_LogitFiltered**

**TableS9_SignalP_prediction_results**

**TableS10_Key_resources**

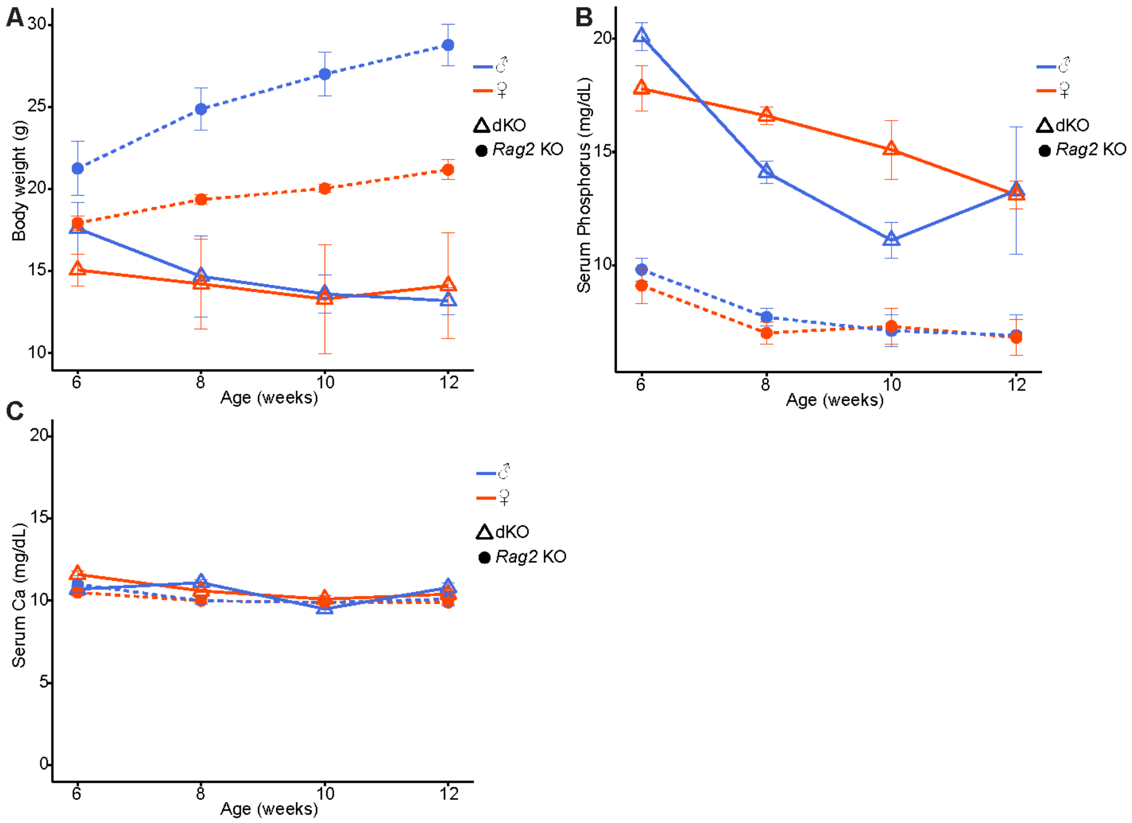


**Supplementary Figure 1. Longitudinal tracking of body weight and systemic mineral metabolism parameters in *Rag2* KO and dKO mice.**

**(A)** Longitudinal measurement of body weight (grams) in male (blue) and female (red) *Rag2* KO (dashed lines, filled circles) and *Rag2* KO × *αKl* KO (dKO; solid lines, open triangles) mice from 6 to 12 weeks of age ($n = 5$ per group, except for male dKO $n = 3$ due to the death of two mice after 9w-old). Both male and female dKO mice exhibited severe growth retardation and age-dependent weight loss, which are typical characteristics of *α-Klotho* deficiency. Data are presented as mean ± SEM.

**(B)** Longitudinal serum phosphorus concentrations (mg/dL) in male and female *Rag2*

KO and dKO mice across 6–12 weeks of age (*n* = 5 per group, except for male dKO *n* = 3 due to the death of two mice after 9w-old). While dKO mice exhibit marked hyperphosphatemia relative to *Rag2* KO controls, serum phosphorus levels do not account for exaggerated male mortality. Data represent mean ± SEM.

**(C)** Longitudinal serum calcium concentration (mg/dL) in male and female *Rag2* KO and dKO mice across 6 to 12 weeks of age (*n* = 5 per group, except for male dKO *n* = 3 due to the death of two mice after 9w-old). Serum calcium levels remained stable and equivalent across all genotypes and sexes. Data represent mean ± SEM.

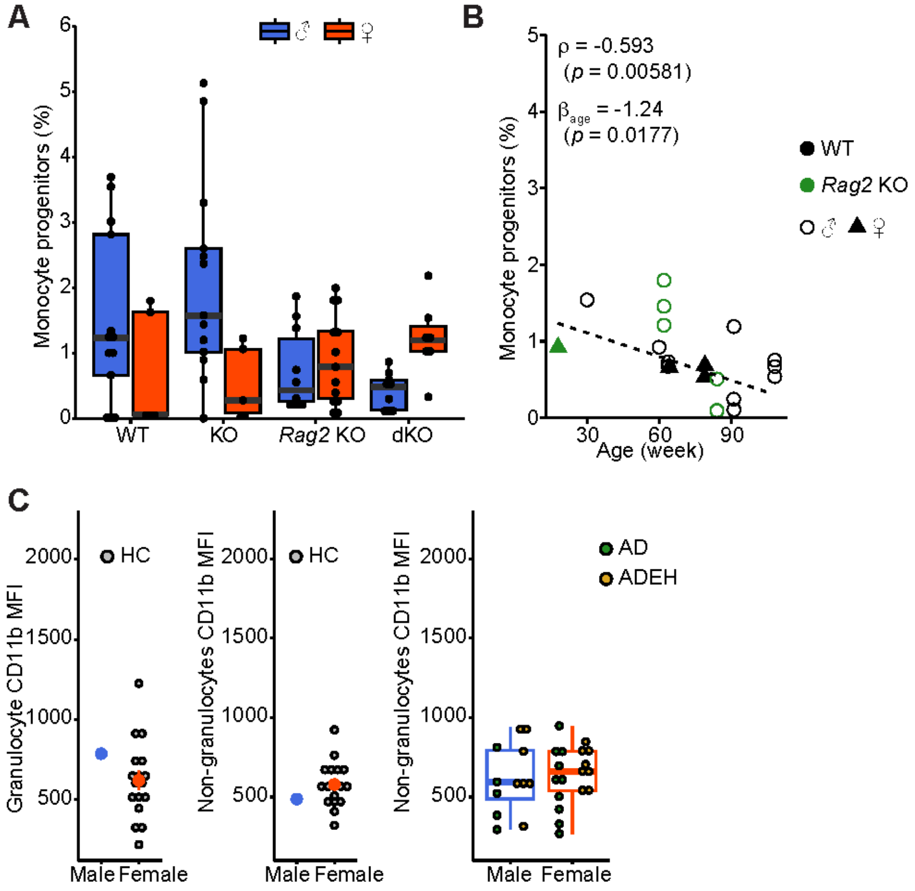


**Supplementary Figure 2. Characterization of non-neutrophils in mouse bone marrow and cell-type specificity of CD11b expression in human cohorts.**

**(A)** Flow cytometric quantification of bone marrow monocyte progenitor frequencies (%) in male (blue) and female (red) WT, *α-Klotho* KO (KO), *Rag2* KO, and double knockout (dKO) mice. Box plots depict the median, interquartile range, and individual biological replicates. Monocyte progenitor proportions showed no consistent sex- or genotype-dependent shifts.

**(B)** Longitudinal correlation between bone marrow monocyte progenitor frequency (%) and age (weeks) in WT (black) and *Rag2* KO (green) mice aged > 12 weeks. Males are indicated by open circles (○), and females by filled triangles (▲). The dashed line represents a linear regression fit. Statistical evaluation demonstrated an age-dependent decline in monocyte progenitors, as assessed by Spearman's rank correlation ($\rho$ = -0.593, $p = 0.00581$) and linear regression ($\beta_{age} = -1.24$, $p = 0.0177$).

**(C)** Flow cytometric quantification of surface CD11b mean fluorescence intensity (MFI) on human circulating immune cell subsets. Left and middle subpanels: Baseline CD11b MFI on circulating granulocytes (left) and non-granulocytes (middle) from healthy control (HC) human subjects stratified by male (blue, $n = 1$) and female (red, $n = 17$). Right subpanel: Surface CD11b MFI on circulating non-granulocytes from human patient cohorts with atopic dermatitis (AD, green circles) or AD complicated by eczema herpeticum (ADEH, yellow circles). Box plots represent the median and interquartile range. In contrast to granulocytes (Figure 2D), non-granulocytes exhibit no significant sex-dependent dimorphism in surface CD11b expression.

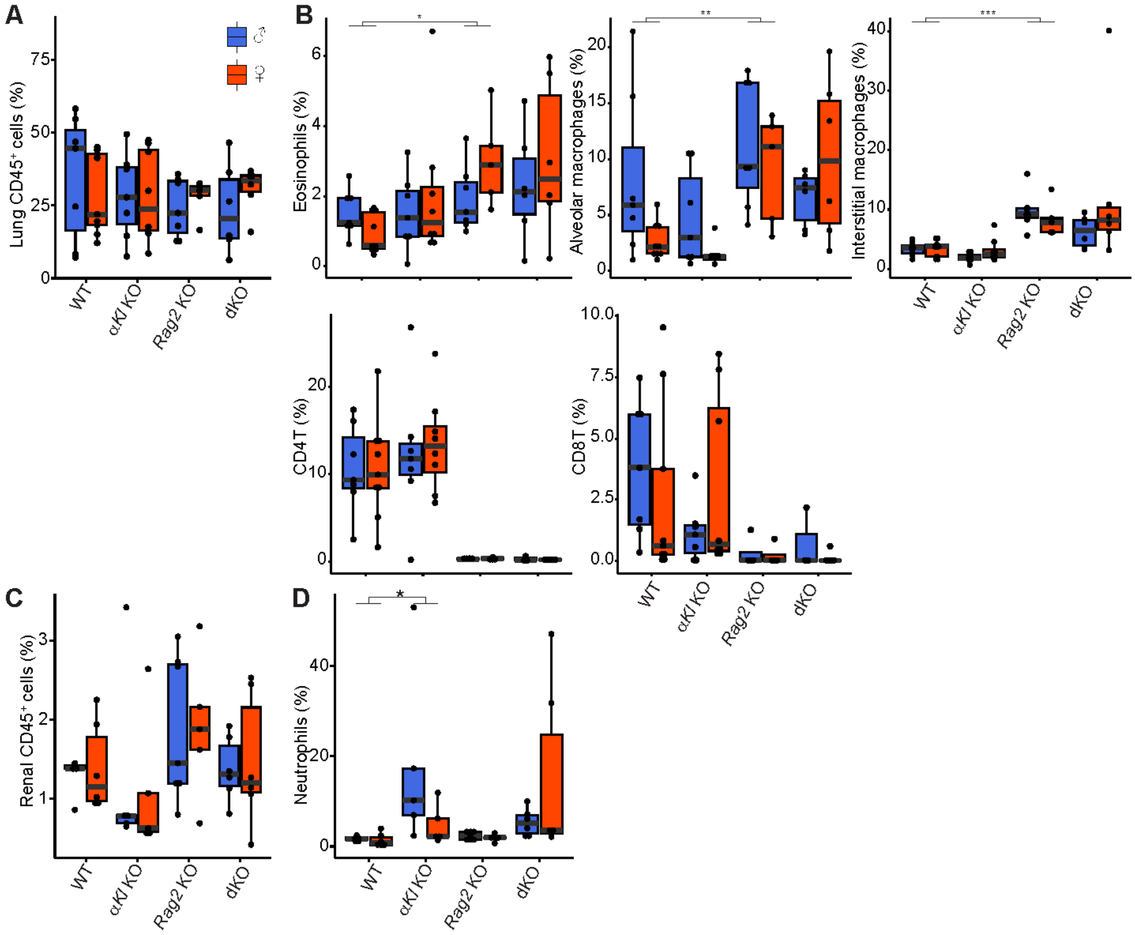


**Supplementary Figure 3. Immune cell profiling in lung and kidney tissues across different genetic backgrounds and sexes.**

**(A)** Flow cytometric quantification of total leukocyte frequencies ($CD45^{+}$ cells, % of total live lung single cells) in lung tissue from male (blue) and female (red) WT, *α-Klotho* KO (*αKl* KO), *Rag2* KO, and double knockout (dKO) mice. Box plots depict the median, interquartile range, and individual biological replicates. Overall, immune cell infiltration into the lungs remained stable across sexes and genotypes.

**(B)** Frequencies of specific non-neutrophil leukocyte subsets in lung tissue expressed as a percentage of total lung $CD45^+$ leukocytes: eosinophils ($SiglecF^+CD11c^-$; top left), alveolar macrophages ($SiglecF^+CD11c^+$; top middle), interstitial macrophages ($Ly6G^-$ $CD11b^+F4/80^+$; top right), $CD4^+$ T cells (bottom left), and $CD8^+$ T cells (bottom right). Expected baseline depletion of $CD4^+$ and $CD8^+$ T lymphocytes is observed in *Rag2*-deficient strains (*Rag2* KO and dKO). Data for B cells are not shown because scarcely any cell was detected in the lung regardless of the *Rag2* genotype. Statistical evaluation by two-way ANOVA followed by post-hoc pairwise comparisons with Benjamin-Hochberg adjustment (*: $p < 0.05$, **: $p < 0.01$, ***: $p < 0.001$ for genotype main effects); no significant sex-dependent differences were detected within any subset.

**(C)** Flow cytometric quantification of total $CD45^+$ leukocyte frequencies (%) in the kidney tissue of male and female mice across genotypes, showing minimal renal immune infiltration.

**(D)** Percentage of neutrophils within total renal $CD45^+$ leukocytes across genotypes and sexes (*: $p < 0.05$ for genotype main effect). In contrast to the robust male-biased neutrophil recruitment in the lung, renal tissue displayed no sex-dimorphic neutrophil accumulation.

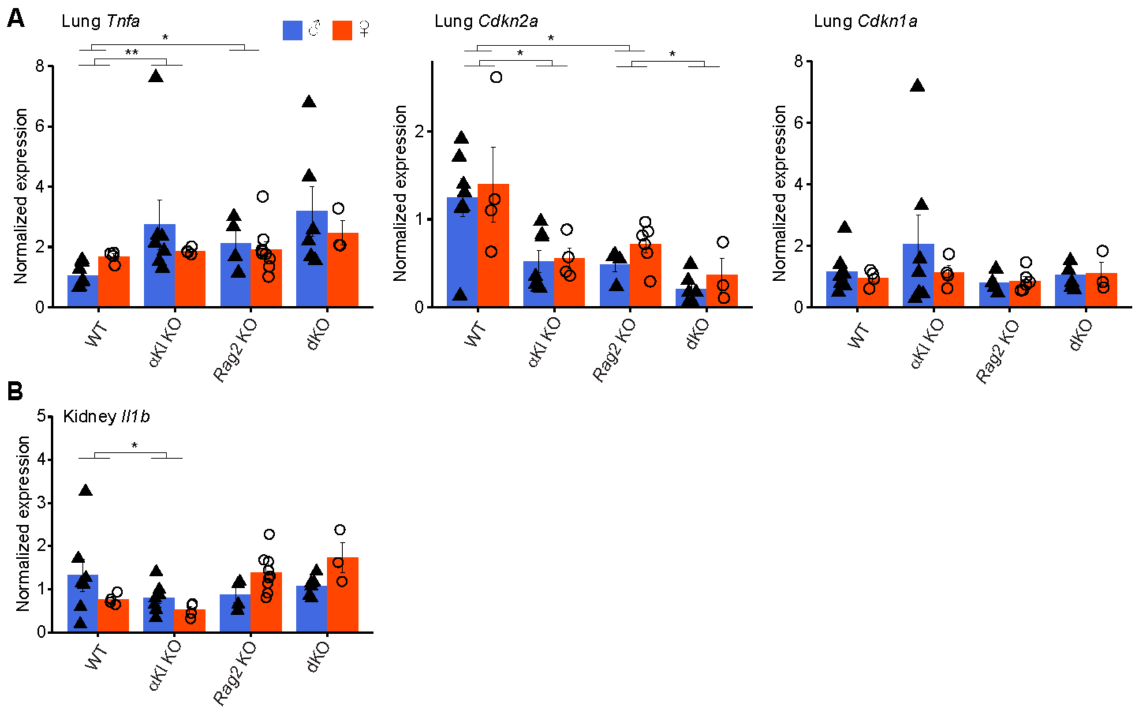


**Supplementary Figure 4. Expression analysis of inflammatory and senescence markers in the lung tissue and *Il1b* transcript levels in the kidney tissue.**

**(A)** Quantitative RT-qPCR assessment of senescence-associated secretory phenotype (SASP) and cellular senescence markers in the whole lung tissue from male (blue, filled triangles) and female (red, open circles) WT, *α-Klotho* KO (*αKl* KO), *Rag2* KO, and double-knockout (dKO) mice: *Tnfa* (left), *Cdkn2a* ($p16^{INK4a}$, middle), and *Cdkn1a* ($p21^{Cip1/Waf1}$, right). Expression levels were normalized to *Actb*. Data represent mean ± SEM with individual biological replicates shown. Data for *Il6* are not shown because scarcely any expression was detected regardless of genotype. Statistical evaluation by two-way ANOVA followed by post-hoc pairwise comparisons with Benjamin-Hochberg adjustment (*: $p < 0.05$, **: $p < 0.01$ for genotype main effects). No significant sex-dependent differences were detected across genotypes for these pulmonary markers.

**(B)** Quantitative RT-PCR analysis of *Il1b* transcript levels in whole kidney tissue across male and female WT, *αKl* KO, *Rag2* KO, and dKO mice (*: $p < 0.05$ for genotype main effect). In contrast to the pronounced male-biased elevation of *Il1b* expression in the lungs (Figure 4A), renal *Il1b* expression displayed no sex-specific accumulation in mutant animals.

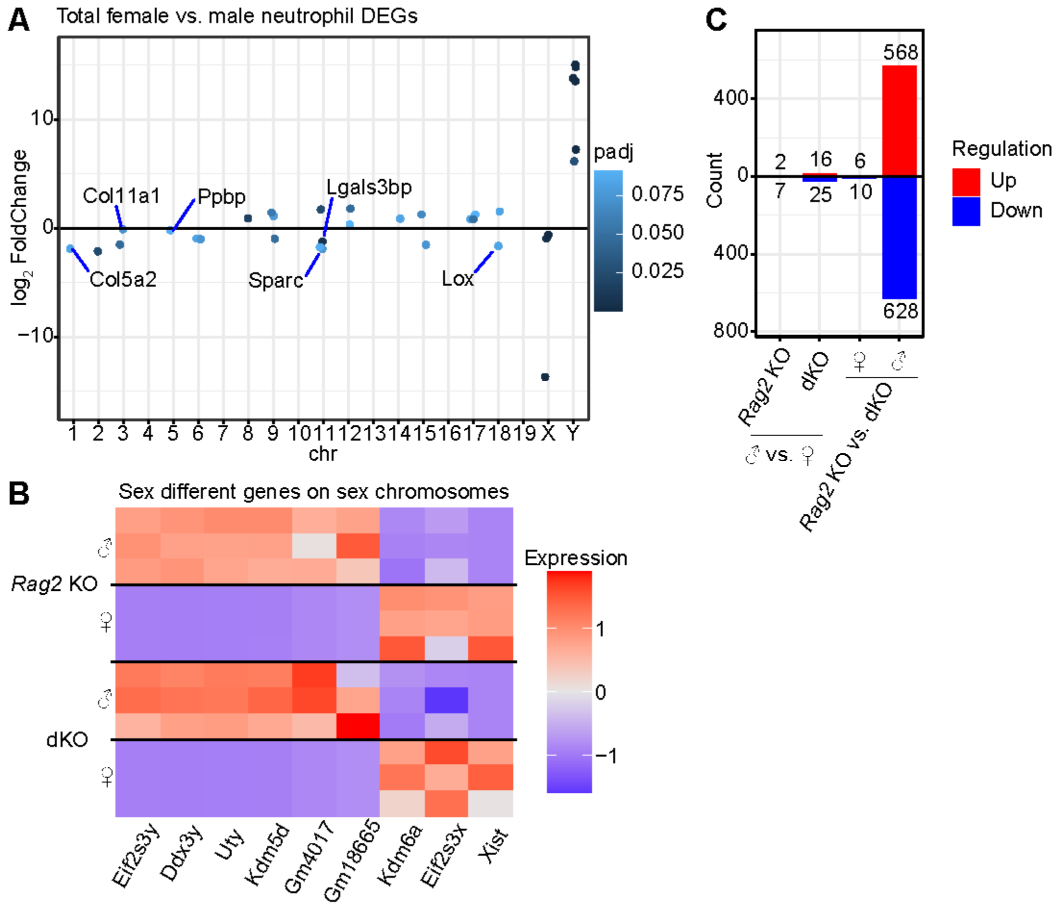


**Supplementary Figure 5. Chromosomal allocation, sex chromosome transcript validation, and differential gene expression counts in bone marrow neutrophils.** **(A)** Chromosomal plot showing the $\log_2$ fold change and genomic distribution of differentially expressed genes (DEGs) comparing female versus male bone marrow neutrophils from both *Rag2* KO and dKO mice across autosomes (chromosomes 1–19) and sex chromosomes (X, Y). The dot color intensity corresponds to the adjusted *p*-value (padj). Key extracellular matrix (ECM) and adhesion-associated autosomal DEGs are explicitly labeled (*Col5a2*, *Col11a1*, *Ppbp*, *Sparc*, *Lgals3bp*, and *Lox*).

**(B)** Expression heatmap (*z*-score normalized) of sex-differentiated genes located on sex chromosomes across male and female *Rag2* KO and double knockout (dKO) neutrophils (*n* = 3 biological replicates per group). Canonical Y-linked transcripts (*Eif2s3y*, *Ddx3y*, *Uty*, *Kdm5d*, *Gm4017*, and *Gm18665*) showed selective expression in males, whereas X-linked transcripts (*Kdm6a*, *Eif2s3x*, and *Xist*) displayed elevated expression in females, confirming sample identity and transcriptomic fidelity.

**(C)** Quantification of significantly upregulated (red) and downregulated (blue) DEGs (padj < 0.1) across pairwise sex (in *Rag2* KO and dKO) and genotype (*Rag2* KO vs. dKO in females and males) comparisons. The numbers above and below the bars indicate the total DEG counts. Note the pronounced transcriptomic remodeling specifically triggered in male neutrophils following *α-Klotho* deficiency (*Rag2* KO and dKO males: 568 upregulated and 628 downregulated genes).

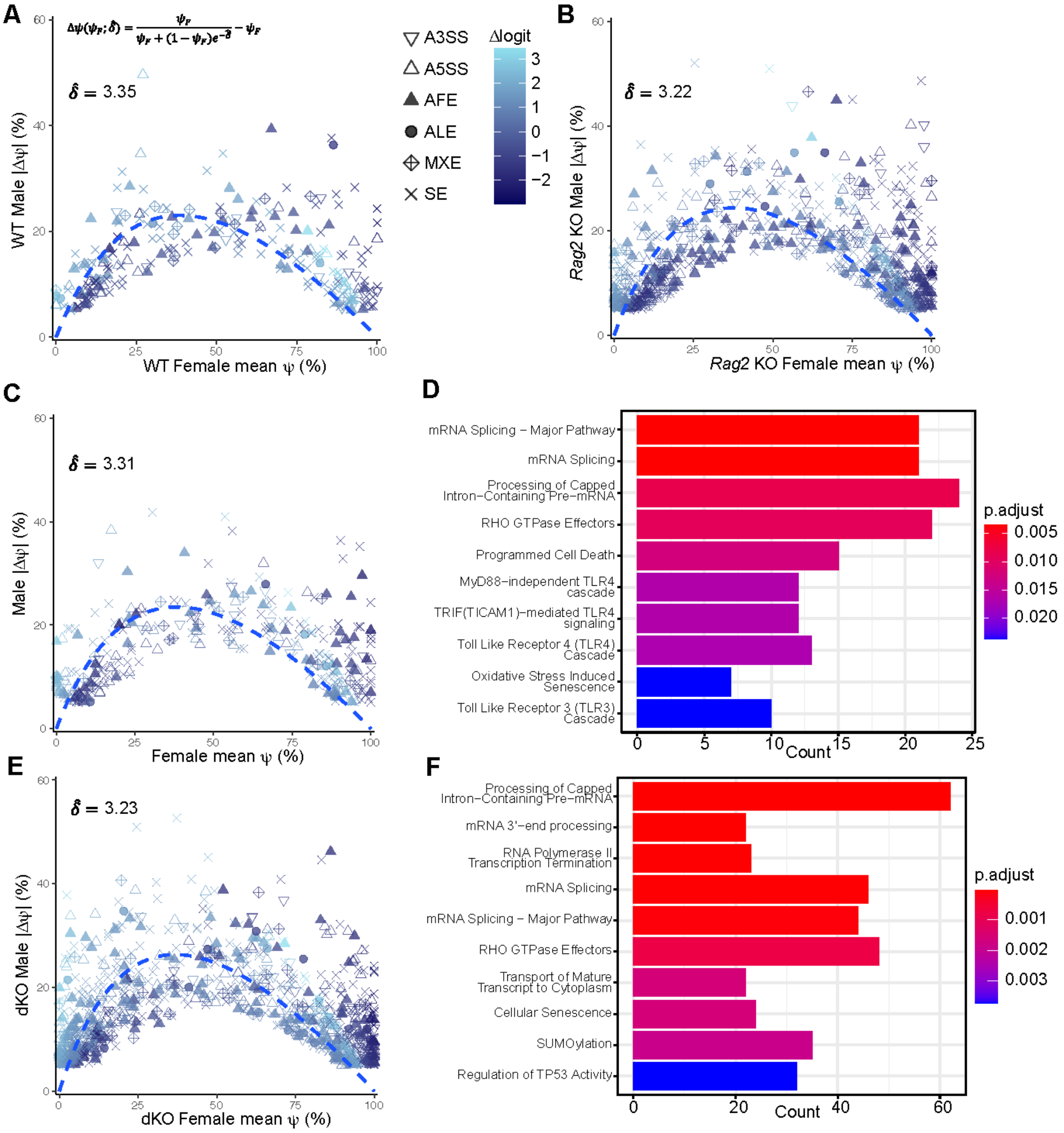


**Supplementary Figure 6. Sex-biased alternative splicing landscapes in wild-type, *Rag2*-deficient, and *Rag2*/*αKl*-double-knockout bone marrow neutrophils.**

**(A)** Distribution of sex-biased alternative splicing (AS) events in young wild-type (WT) bone marrow neutrophils re-analyzed from public RNA-seq datasets, plotting male Percent Spliced In difference (|Δψ|, %) against female mean ψ (%) ($\hat{\delta}$ = 3.32). The categorized splicing event types include alternative 3' splice sites (A3SS), alternative 5' splice sites (A5SS), alternative first exons (AFE), alternative last exons (ALE), mutually exclusive exons (MXE), and skipped exons (SE). The color intensity of the points corresponds to the change in log odds (Δlogit).

**(B)** Alternative splicing landscape in bone marrow neutrophils from *Rag2* KO mice (own RNA-seq dataset), plotting male |Δψ| (%) versus female mean ψ (%), with events colored by change in log-odds (Δlogit) ($\hat{\delta}$ = 3.22).

**(C)** Distribution of sex-biased alternative splicing (AS) events in bone marrow neutrophils from 20-month-old WT mice re-analyzed from public RNA-seq datasets, plotting male |Δψ| (%) against female mean ψ (%) ($\hat{\delta}$ = 3.29). The color intensity of the points corresponds to the change in log odds (Δlogit).

**(D)** Reactome pathway enrichment analysis of differentially spliced transcripts in 20-month-old WT male versus female neutrophils. The bar length indicates the gene count, and the fill color corresponds to the adjusted *p*-value (p.adjust). Prominent enrichment was observed in core mRNA splicing and processing machinery, RHO GTPase effectors, programmed cell death, and Toll-like receptor signaling cascades (TLR4, TRIF-mediated, MyD88-independent, and TLR3).

**(E)** Alternative splicing landscape in bone marrow neutrophils isolated from double-knockout (dKO) mice (own RNA-seq dataset), plotting male |Δψ| (%) versus female mean ψ (%) ($\hat{\delta}$ = 3.26). Events are colored according to changes in log odds (Δlogit).

**(F)** Reactome pathway enrichment analysis of differentially spliced gene sets in dKO female versus male neutrophils. The bar length indicates the gene count, and the fill color corresponds to the adjusted *p*-value (p.adjust). Prominent enrichment was observed in pre-mRNA processing, splicing machinery, RHO GTPase effectors, cellular senescence, and SUMOylation pathways.
